# NAD(H)-mediated tetramerization controls the activity of *Legionella pneumophila* phospholipase PlaB

**DOI:** 10.1101/2020.09.01.246603

**Authors:** Maurice Diwo, Wiebke Michel, Philipp Aurass, Katja Kuhle-Keindorf, Jan Pippel, Joern Krausze, Christina Lang, Wulf Blankenfeldt, Antje Flieger

## Abstract

The virulence factor and phospholipase PlaB promotes lung colonization, tissue destruction, and intracellular replication of *Legionella pneumophila*, the causative agent of Legionnaires’ disease. It is exposed at the bacterial surface and shows an extraordinary activation mechanism by tetramer deoligomerization. To unravel the molecular basis for enzyme activation and localization, we determined the crystal structure of PlaB in its tetrameric form. We found that the tetramer is a dimer of identical dimers, and a monomer consists of an N-terminal phospholipase α/β-hydrolase domain augmented by two non-canonical two-stranded β-sheets, β6/β7 and β9/β10. The C-terminal domain reveals a novel fold displaying a bilobed β-sandwich with a hook structure that is required for dimer formation and complementation of the phospholipase domain in the neighboring monomer. Unexpectedly, we observed eight NAD(H) molecules at the dimer/dimer interface, suggesting that these molecules stabilize the tetramer and hence lead to enzyme inactivation. Indeed, addition of NAD(H) increased the fraction of the tetrameric form and concomitantly reduced activity. β9/β10 mutants revealed a decrease in the tetrameric fraction, altered activity profiles, and mislocalization. Protein variants lacking the hook or strands β6/β7 were unaffected in terms of localization but lost their activity, and lid mutants changed substrate specificity. Together, these data reveal structural elements and an unprecedented NAD(H)- mediated tetramerization mechanism required for spatial and enzymatic control of a phospholipase virulence factor. The regulatory process identified is ideally suited to fine tune PlaB in a way that protects *L. pneumophila* from self-inflicted lysis while ensuring its activity at the pathogen–host interface.

## Introduction

Phospholipases are important enzymes in infectious disease pathogenesis involved in host modulation and damage. They have been assigned to different groups depending on the preferred cleavage site within the phospholipid substrate. Phospholipases A (PLAs) and lysophospholipases A (LPLA) hydrolyze carboxyl ester bonds at the sn-1 or sn-2 position in phospholipids or lysophospholipids, respectively, and release fatty acids (1, 2). In *Legionella pneumophila*, a Gram-negative bacterium that causes Legionnaires’ disease, at least

15 genes encoding PLAs/LPLAs belonging to three families are found. Many of these are secreted to modulate the host cell but only one, PlaB, is uniquely presented at the bacterial surface (1, 3–6). In this study, we focused on PlaB, a hemolysin and virulence factor that promotes intracellular replication in macrophages by its PLA and LPLA, activities (7–9). PlaB is also crucial for lung colonization and tissue destruction in guinea pig infections (6). It is the only characterized member of a recently discovered PLA family, and homologs are found in several water-associated bacteria including the opportunistic pathogen *Pseudomonas aeruginosa* (7, 8). Previous work suggested that PlaB is organized into two domains, namely an N-terminal phospholipase (amino acids 1-∼300) connected to a C-terminal domain (CTD) (amino acids ∼301-474) that is also essential for activity (8). The catalytic triad S85/D203/H251 of the N-terminal domain of PlaB and its homologs is embedded in uncommon consensus motifs which are unique among lipases (8). The CTD is not related to formerly characterized proteins, but we have earlier shown that the last 15 amino acids of PlaB are necessary for activity, although their exact role is not understood (8, 9). Subcellular fractionation and proteinase K digests revealed that PlaB is associated with the outer membrane (OM) and exposed on the surface. However, due to the apparent lack of export signal sequences, lipid anchors, or transmembrane helices, the determinants for export and membrane-association remain elusive (6, 7).

Since PlaB represents the most active PLA/LPLA in *L. pneumophila*, affecting lipids, such as phosphatidylcholine (PC) and phosphatidylglycerol (PG), found in the lung of the human host and in *Legionella* (7, 10–12), fine-tuned control of enzyme activity may be crucial to prevent damage to the pathogen itself. Indeed, PlaB shows an extraordinary activation mechanism that requires protein deoligomerization. At higher PlaB concentrations, the enzyme occurs in an inactive tetrameric form, whereas it deoligomerizes in the lower nanomolar concentration range where it possesses its highest specific activity (9). However, the mechanism behind enzyme regulation is not understood.

To decipher the molecular basis for PlaB’s unusual activation and to gain insight on how it associates with the OM, we have determined its crystal structure. This allowed us to identify important structural features and, interestingly, revealed an NAD(H)-mediated tetramerization mechanism that controls activity.

## Results

### PlaB crystallizes as a dimer of dimers and a hook structure is required for dimer formation

Extensive efforts were required to obtain high-quality crystals of PlaB. Most crystals suffered from anisotropic diffraction and streaky, overlapping reflections. The structure was finally determined at 2.3 Å resolution by single-wavelength anomalous dispersion of seleno-*L*-methionine-labeled PlaB in a crystal obtained by seeding (Fig. S1A and B). This crystal belonged to space group P2_1_ and displayed strong translational non-crystallographic symmetry, explaining why the refinement converged at very high R-factors for this data set (Tab. S1). The asymmetric unit comprised four chains that arrange in a tetrameric fashion, best described as a dimer of dimers (Fig. 1A). Interestingly, the contact area between both dimers is relatively small and PISA analysis of the protein chains alone did not recognize this tetramer as a stable assembly (13). The PlaB dimers, however, are predicted as stabilized by approximately - 29.8 kcal/mol. The dimers are formed by head-to-tail interaction between the N-terminal phospholipase domain of the first monomer and a protruding hook-like extension at the C-terminus of the second (residues R446-D474) (Fig. 1A, B and C, Tab. S2). This leads a) to an extension of the central β-sheet of the phospholipase domain by β18 from the hook region, explaining the previously determined loss of activity when residues from the C-terminus are deleted (8, 9), and b) to a number of polar and hydrophobic interactions between the last ∼30 amino acids and residues of the N-terminus.

**Figure 1.**
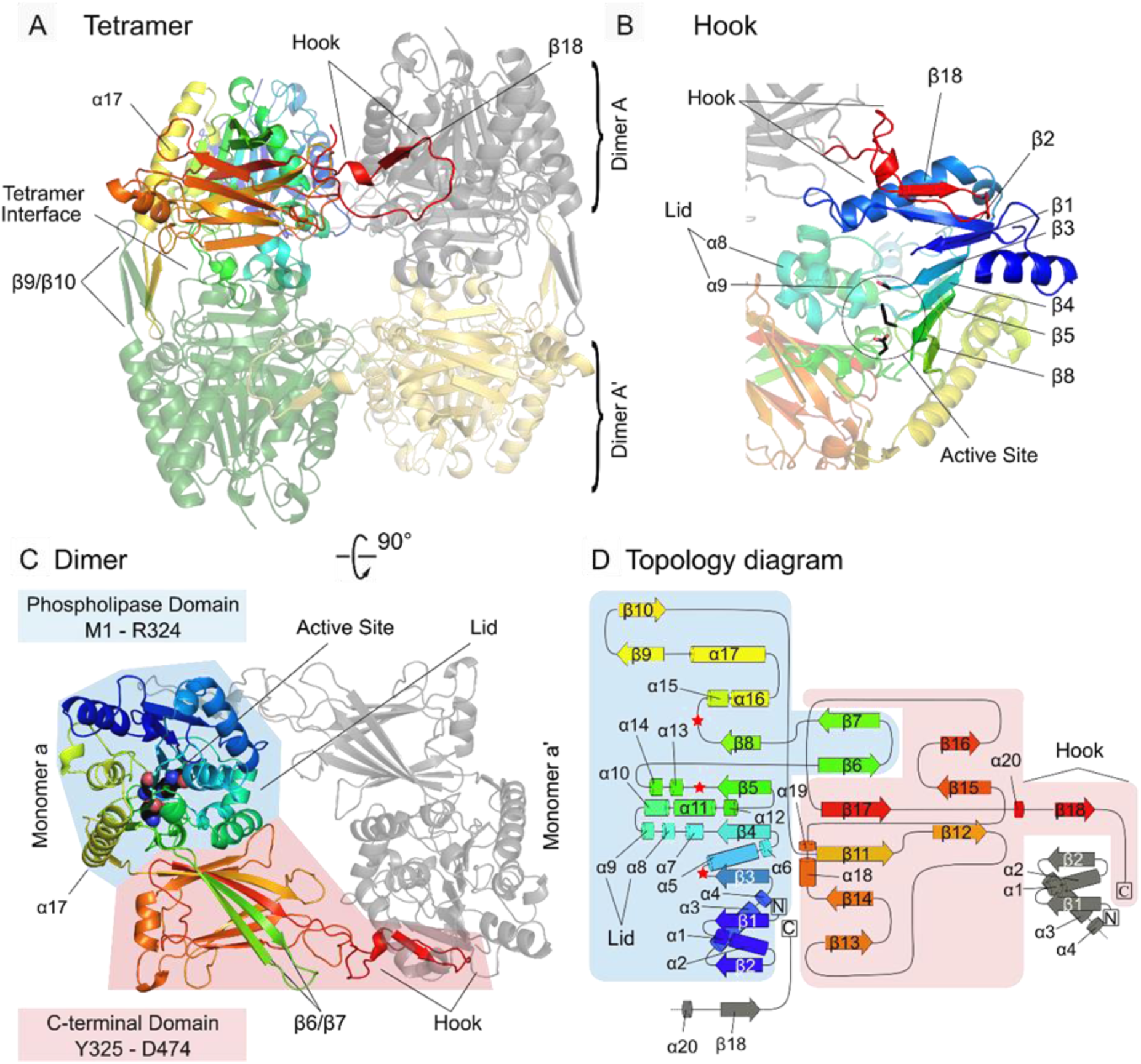
PlaB crystallizes as a dimer of dimers and a hook structure is required for dimer formation. A) The PlaB tetramer is built from two identical dimers: dimer A (color gradient and grey) and dimer A’ (green and yellow). α17 leads to β9/β10 which participate in the tetramer interface. The CTD comprises a hook structure connecting two PlaB molecules by a head to tail interaction. B) β18 located in the hook (shown in red) complements the internal β-sheet (β2/β1/β3/β4/β5/β8 in the order of structural succession) from the α/β-hydrolase fold by attaching to β2 in the NTD of the second PlaB molecule of the dimer. The active site is covered by a lid in the closed conformation (α8/α9). C) The PlaB dimer consists of an N-terminal phospholipase domain (blue background) and a C-terminal novel bilobed β-sandwich domain (red background). Dimer A is shown (color gradient and grey for the two monomers). The active site (S85/D203/H251) is buried inside the ABH-fold of the NTD and is covered by a lid structure in the closed conformation. The β6/β7-sheet protrudes from the NTD into the CTD. The hook forms the dimer interface, complementing the α/β-hydrolase fold of a second PlaB molecule. D) Topology diagram of PlaB (color gradient). Background coloring refers to the division of PlaB into the NTD and CTD. The location of the active site is indicated (red stars). The β-sheet of the N-terminal ABH-fold is assembled from the parallel strands β2, β1, β3, β4, β5 and β8 and antiparallelly complemented by β18 from the second PlaB molecule of the dimer. The CTD has a mixed configuration of β-strands. While β7, β6, β17, β11, β14 and β13 form the upper lobe, β16, β15 and β12 form the lower lobe of a bilobed β-sandwich.

### The phospholipase domain is a typical α/β-hydrolase extended by non-canonical elements

The PlaB monomer consists of two clearly distinguishable domains (Fig. 1C and D). These comprise residues 1-∼324 and residues ∼325-474, representing the phospholipase domain and a CTD consisting of a unique bilobed β-sandwich extended by the hook structure, respectively. The N-terminal domain (NTD) displays a typical α/β-hydrolase (ABH) fold, hallmarked by a central six-stranded parallel β-sheet (spatially ordered as β2-β1-β3-β4-β5-β8) that is surrounded by 17 α- helices (α1 to α17) (Fig. 1A, C and D) (14, 15). The catalytic center is readily discernible by the catalytic triad S85/D203/H251, which was previously identified through sequence alignments and mutagenesis (Fig. 1B, C) (8). It is shielded from the solvent by a closed lid formed by residues G127-G144 of α8 and α9, as is frequently observed in ABHs in the absence of substrates (Fig. 1B, Fig. 2A and B, Tab. S2) (16). The electron density indicates that this lid is highly flexible. In addition to these canonical features, the NTD also possesses unique structural elements that are specific to PlaB, namely a long α-helix (α17, residues R285 - N305) and two anti-parallel β-sheets (β6/β7, residues S216-R237 and β9/β10, residues K308-I321) (Fig. 1A, C and D, Tab. S2). While the α-helix participates in lining the large central β-sheet, the additional β-sheets β6/β7 and β9/β10 protrude from the ABH (Fig. 1A, and 1C).

**Figure 2.**
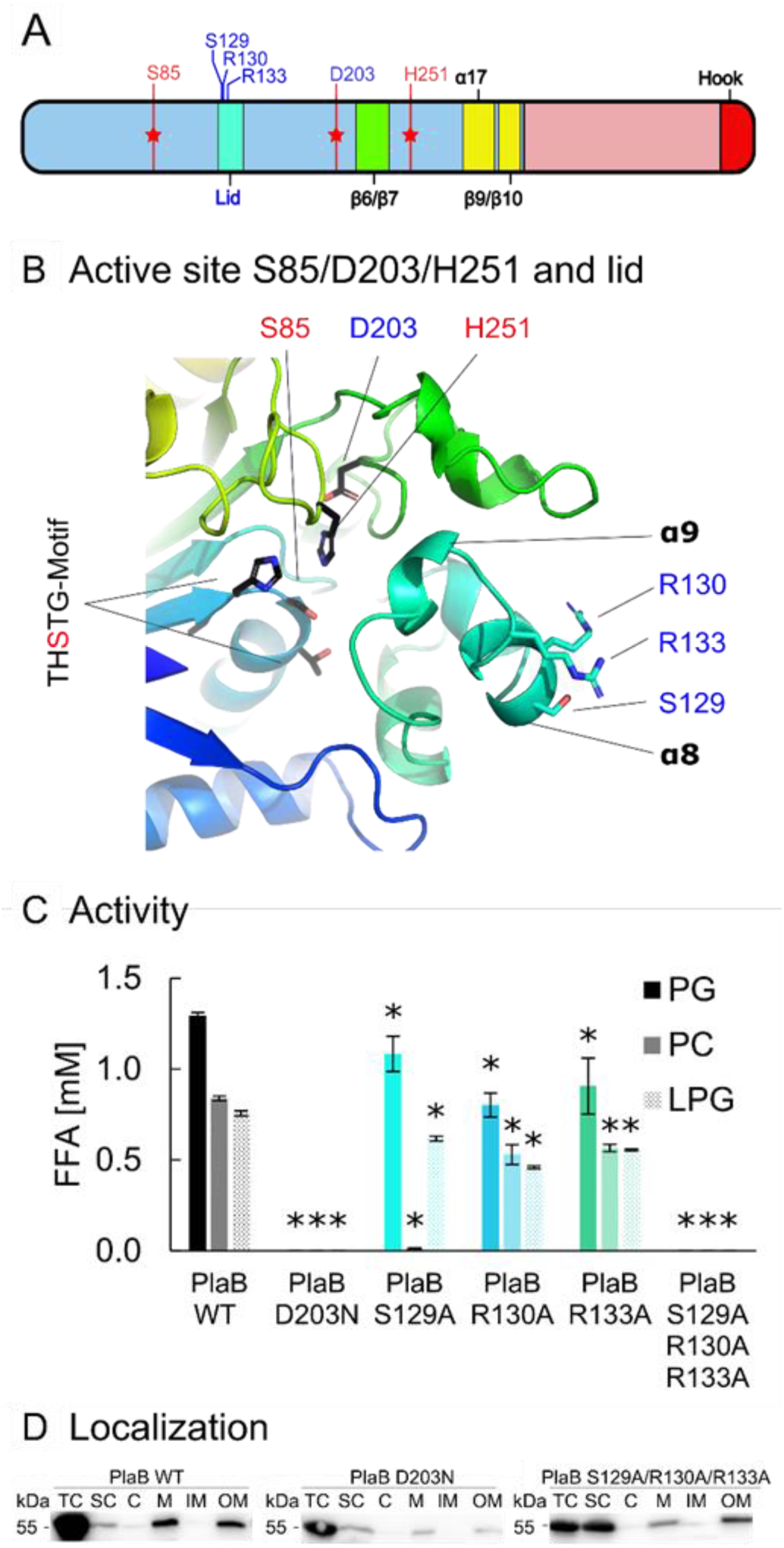
The lid structure of PlaB influences substrate specificity but not localization. A) Overview of PlaB major sequence and structural features, such as the active site (red stars) and residues mutated in the context of this figure (blue text). B) Active site (S85/D203/H251) and lid (α8/α9, cyan) in their structural context. S85 is embedded in the THSTG motif. Residues that have been probed by mutagenesis are indicated by blue text. C) Enzymatic activity of PlaB WT (black) and mutant strains (D203N, no activity; S129A, cyan; R130A, blue; R133A, green; and S129A/R130A/R133A, no activity) towards different phospholipid substrates. 1:100 diluted cell lysates of *E. coli* expressing different PlaB versions were incubated with the different lipids for 15 min and subsequently quantities of released fatty acids (FFA) were determined. Error bars indicate SD. Statistical analysis was performed using two-tailed, unpaired Student’s t-tests, relating PlaB WT to PlaB versions. * p < 0.02 (N=3). Western blot of cell lysates, see Fig. S4. Localization of different PlaB versions. Western blot analysis after cell fractionation of *E. coli* expressing different PlaB versions using an anti-strep-tag antibody. Please refer to Fig. S3 for PlaB WT or D203N mutant analysis including fractionation controls. Abbreviations: TC – total cell lysate, SC – soluble content, C - cytosol, M – membrane, IM – inner membrane, OM – outer membrane.

### The C-terminal domain possesses a novel bilobed β-sandwich structure

The CTD of PlaB is dominated by a bilobed β-sandwich consisting of a six-stranded mixed (β6- β7-β11-β13-β14-β17) and a three-stranded anti-parallel β-sheet (β12-β15-β16) (Fig. 1C and D). Interestingly, two strands of the first β-sheet stem from the NTD (β6-β7), suggesting that the CTD requires the phospholipase domain for stability (Fig. 1C and D). Searches with DALI (17) did not identify any similar structures, indicating that the PlaB CTD constitutes a novel fold.

### Lid residues are central to activity and substrate specificity

Before analyzing the importance of residues and structural motifs for enzyme activity and localization, we confirmed membrane, OM, and surface localization of PlaB in *L. pneumophila* by means of a) subcellular fractionation (Fig. S2A and B), b) proteinase K digestion (Fig. S2C), and surface-protein biotin labeling (Fig. S2D). Since PlaB sequence and previous results did not suggest involvement of export signals and modes characterized in *L. pneumophila* for PlaB export (6), we tested its activity and localization after gene expression in *E. coli*. We found that PlaB showed activity and most interestingly localized to the OM (Fig. 2C, 2D, and S3). The catalytic triad mutant D203N, devoid of activity, localized to the OM likewise (Fig. 2C, 2D, and S3).

A lid is a typical feature of lipases regulating substrate access to the catalytic site. Both its amphipathic nature and specific amino acids are crucial for activity and specificity (18). Indeed, we found that the lid point mutation S129A revealed severely reduced activity towards PC, a phospholipid with a positive-charged head group, but only slightly reduced activity towards negative-charged PG and lysophosphatidylglycerol (LPG) (Fig. 2B and C). This is in accordance to previous findings showing that this residue, now recognized as part of the lid, promotes PC hydrolysis and hemolysis (8). Further, the positive charge of the neighboring lid residues R130 and R133 could contribute to the recognition of negatively charged lipid substrates, as has been observed for other lid-containing hydrolases (14, 16). Whereas point mutations led to reduced activity, towards PG and LPG, but also influenced PC hydrolysis, the triple mutant S129A/R130A/R133A was inactive towards all substrates tested (Fig. 2C). Localization of the latter protein variant at the OM was not affected (Fig. 2D). These findings demonstrate that the lid region influences activity and substrate specificity.

### The non-canonical β-strand β6/β7 and the hook are essential for enzyme activity

The non-canonical β-strands β6/β7 and β9/β10 are anti-parallel to each other and project away from the ABH fold (Fig. 1A and C). Given the importance of strands β6 and β7 within the CTD, it is not surprising that the respective deletion mutant was inactive, yet it still localized to the OM (Fig. S5A, B, and C). A similar phenotype, i.e. loss of activity and unchanged localization, was found when the hook (residues R446-D474) was deleted (Fig. S5A, B and C). This is in accordance with previous observations where shortening of the CTD led to a decrease in activity (9), corroborating our finding that the hook connects two PlaB monomers and extends the ABH fold to yield active PlaB dimers.

### β9/β10 influences enzyme activity and contributes to OM localization of PlaB and tetramer stability

The β9/β10 deletion mutant lost its activity against PC but was still active towards PG and LPG, although at a lower level. Therefore, β9/β10 affects the substrate spectrum of PlaB (Fig. 3A, B, C). Interestingly, the mutant showed subcellular mislocalization and was associated with the inner membrane and the culture supernatant, suggesting that OM association depends on β9/β10 (Fig. 3D and E). Furthermore, we analyzed truncations of PlaB by means of proteinase K digests and found that amino acids 302-323 contributed to surface presentation (Fig. S6) corroborating the importance of β9/β10 (K308-I321) (Table S2). β9/β10 reveals an intriguing pattern of hydrophobic and cationic residues displayed on both sides of the sheet (K308, F310, K312, F316, R318, Y320) (Fig. 3B). Similar motifs contribute to cation-π interactions in proteins that bind to positively-charged phospholipid head groups (19, 20), and the corresponding triple mutant F310D/F316D/Y320D indeed resembled the β9/β10 deletion mutant in terms of activity and localization (Fig. 3C, D and E).

**Figure 3.**
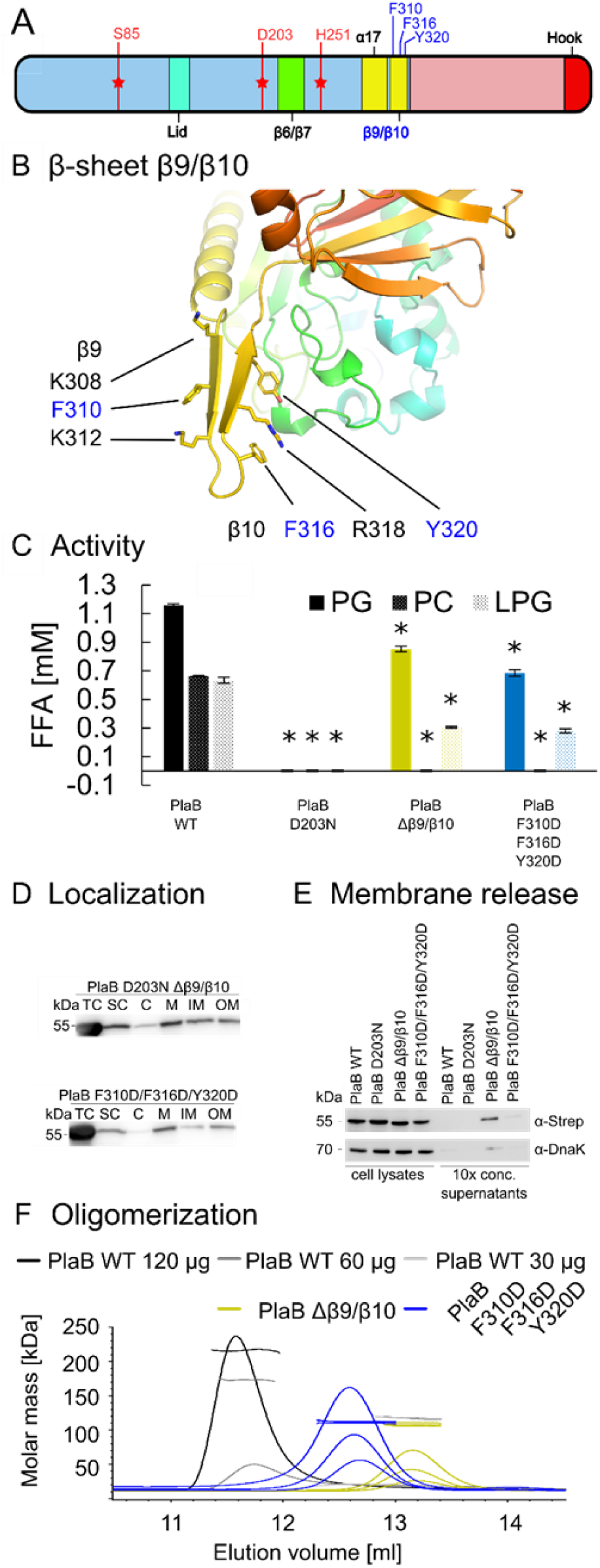
β9/β10 of PlaB affects activity, localization, and tetramer stability. A) Overview of PlaB major sequence and structural features, such as the active site (red stars) and residues mutated in the context of this figure (blue text). B) β9/β10 (yellow) protrudes from the NTD and is decorated with cationic and aromatic residues. Mutated residues are shown in blue. C) Enzymatic activity of PlaB WT (black) and mutants (D203N, no activity; Δβ9/β10, yellow; and F310D/F316D/Y320D, blue) towards different phospholipid substrates. 0.233 nM of different PlaB versions were incubated with the indicated lipids for 30 min and subsequently quantities of released fatty acids (FFA) were determined. Error bars indicate SD. Statistical analysis was performed using two-tailed, unpaired Student’s t-tests, relating PlaB WT to PlaB versions. p < 0.02 (N=3). SDS PAGE analysis of proteins, see Fig. S4. D) Localization of different PlaB versions. Western blot analysis after cell fractionation of *E. coli* expressing different PlaB versions using an anti-strep-tag antibody. Please refer to Fig. 2D or S3 for PlaB WT or D203N mutant analysis and to Fig. S7 for fractionation controls. Abbreviations: TC – total cell lysate, SC – soluble content, C - cytosol, M – membrane, IM – inner membrane, OM – outer membrane. Western blot analysis after separation of *E. coli* expressing different PlaB versions into cell lysate and culture supernatant and detection of PlaB using an anti-strep-tag antibody or cytosolic control protein DnaK using a DnaK antibody. F) SEC-MALS analysis of PlaB WT and β9/β10- mutants shows concentration-dependent oligomerisation. Application of 120 μg PlaB WT (black) shows tetramer (1.94 µM), 60 μg mixture of tetramers and dimers (1.14 and 0.34 µM respectively) and 30 μg, dimeric PlaB (0.17 µM. PlaB Δβ9/β10 (yellow) and F310D/F316D/Y320D (blue) were found as dimers (1.02/0.57/0.26 µM and 2.77/1.34/0.81 µM, respectively). Detailed results are shown in Table S3.

As suggested in the representation of the tetramer structure, sheet β9/β10 may be involved in dimer/dimer interactions (Fig. 1A). Therefore, we hypothesized that β9/β10 influences tetramer formation. In fact, as size exclusion chromatography-multiangle light scattering (SEC-MALS) experiments showed, its mutation impeded concentration-dependent tetramerization. Specifically, whereas different concentrations of PlaB wild type revealed either the tetramer only or a tetramer-dimer mixture, the β9/β10 deletion or F310D/F316D/Y320D mutants were present in a dimeric form regardless of protein concentration (Fig. 3F, Table S3). Together these data show, that β9/β10 not only supports the localization of PlaB at the OM and tunes its selectivity towards PC, but also contributes to stabilization of the tetrameric state.

### PlaB phospholipase activity is controlled by NAD(H)-mediated tetramerization

To our surprise, we observed copurified ligands at the tetramer interface, albeit at low occupancy. The shape of their electron density suggested their identity as NAD(H) molecules that bind to two different but closely neighbored binding sites per monomer (Fig. 4A and B). Several experiments revealed that only PlaB tetramers but not the dimers bind NAD(H). In detail, we observed that a) the presence of NAD(H) in the SEC-MALS running buffer led to the exclusive formation of tetramers (Fig. 4C), b) in freshly purified PlaB, the A260/A280 ratio in the SEC-MALS experiment was higher than expected for the peak corresponding to tetrameric PlaB, but not for lower oligomers (Fig. S8),, c) the addition of NAD(H) led to a more defined melting point (Fig. S9), and thio-NAD (SNAD), a component of the thermal shift assay used to optimize the storage buffer for PlaB, improved and accelerated crystallization of the PlaB tetramer, yielding crystals in spacegroup P1 that diffracted up to 1.8 Å and that grew to full size after 20 instead of 150 days (Fig. S1C). The corresponding structure indeed revealed full occupation of the eight NAD(H) binding sites with SNAD molecules (Fig. 4A). Several residues are involved in specific interactions with the ligand, particularly from the NTD (β9/β10, α13, lid) of one PlaB dimer and from the CTD (β12, β13, β14) of the other, also including π-stacking between the nicotinamide group and Y190 and Y196 (Fig. 4B). Interestingly, while all eight NAD(H) binding sites are occupied with SNAD, a substantial part of the lid of all four monomers became extremely flexible such that residues 133-141 (RIKSFFEGI) could not be traced in the corresponding structure. The non-canonical strands β9/β10 of one dimer line the outer NAD(H) binding site, and R318 is involved in a hydrogen bond with the ribose unit of the nicotinamide half of the ligand (Fig. 4B, Fig. S10).

**Figure 4.**
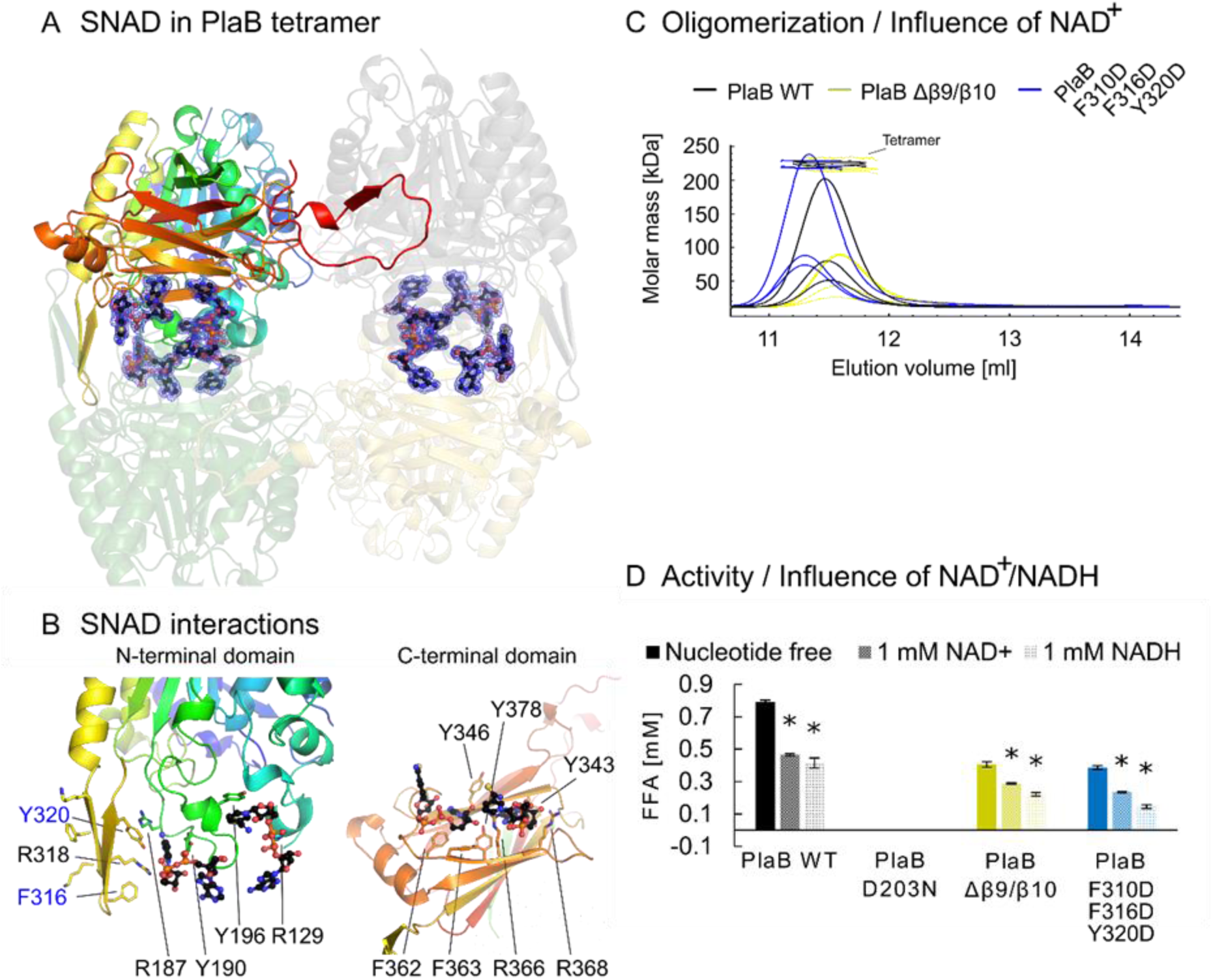
PlaB phospholipase activity is inhibited by NAD(H)-mediated tetramerization. A) The position of eight NAD(H) binding sites within the PlaB tetramer, shown in the same orientation as in Fig. 1A. Note that the identity of these ligands is the thio-derivative SNAD, since the respective data were obtained from a crystal generated in the presence of 1 mM SNAD (Fig. S1C). The blue mesh indicates the final |2F_O_ – F_C_| electron density for the ligands, displayed at 1 σ. B) Two SNAD molecules (black carbons, blue nitrogens, red oxygens, orange phosphorus) per PlaB monomer are coordinated at the tetramer interface. The NTD (left) contributes with β9/β10 (yellow), α13 (green) and the lid structures (cyan) and the CTD (right) with β12, β13 and β14 (orange) to SNAD coordination. Cationic and aromatic side chains (black labels) and mutated residues are shown (blue labels). C) SEC-MALS analysis of PlaB and β9/β10-mutants in the presence of 1 mM NAD^+^ shows tetramers regardless of protein concentration and version (PlaB WT, black, 1.12/0.44/0.24 µM; PlaB Δβ9/β10, yellow, 0.48/0.20/0.09 µM) and PlaB F310D/F316D/Y320D, blue, 1.52/0.51/0.37 µM). Detailed results are shown in Table S3. D) Enzymatic activity of PlaB WT (black) and mutant strains (D203N, no activity; Δβ9/β10, yellow; and F310D/F316D/Y320D, blue) towards PG with/without NAD+ or NADH addition. 0.233 nM of different PlaB versions were incubated with the different lipids for 30 min and subsequently quantities of released fatty acids (FFA) were determined. Error bars indicate SD. Statistical analysis was performed using two-tailed, unpaired Student’s t-tests, relating PlaB versions without nucleotides to PlaB versions supplemented with NAD(H). * p < 0.02 (N=3). SDS PAGE analysis of proteins, see Fig. S4.

As outlined above, PISA analysis did not identify the PlaB tetramer as a stable assembly in the absence of the ligand. We subsequently analyzed the crystal structure of PlaB in the ligand-bound form and found that the tetramer was stabilized by SNAD (ΔG = - 44.7 kcal/mol with SNAD and - 3.0 kcal/mol without SNAD) (13). This suggests that tetramerization and hence inactivation of PlaB is NAD(H)-mediated and we indeed confirmed that NAD^+^ and NADH inhibited the activity of wild type PlaB (Fig. 4D). Although residues of β9/β10 among others were found in close proximity to the ligand, the activity of the β9/β10 deletion mutant and the F310D/F316D/Y320D mutant was also reduced by NAD(H), demonstrating the importance of additional NAD(H) interaction sites (Fig. 4A and 4B). Further, it was possible to reconstitute tetramers for all concentrations of the wild type and mutant proteins when NAD(H) was added. This shows that the tetrameric form of the mutants, although less stable without ligand addition (Fig. 3F), was still stabilized by the ligand (Fig. 4C). We therefore conclude that NAD(H) enhances PlaB tetramerization and concomitantly enzyme inactivity.

### Discussion

Here, we determined the crystal structure of *L. pneumophila* PlaB, demonstrating that it consists of a dimer of dimers. PlaB contains an ABH in its NTD (14), followed by a novel type of β- sandwich domain in the CTD that is completed by two non-canonical β-strands from the NTD (β6/β7). The other additional two-stranded β-sheet β9/β10 of the NTD fulfils several roles for the enzyme. Its deletion indicated requirement for substrate preferences towards PC and its association to the OM. This is probably rooted in patterns of cationic and aromatic amino acids that protrude from β9/β10 and are known to promote phospholipid interaction by means of cation- π interactions (19, 20). The finding of a C-terminal hook-like structure connecting two monomers and contributing a strand to the central ABH β-sheet was unprecedented but explains the importance of C-terminal residues for enzyme activity (9). Similar to many ABH enzymes studied in the absence of substrate (21), tetrameric PlaB contains a flexible lid adopting a closed conformation. Tetramerization sequesters the lids of all four PlaB molecules at the tetramer interface, thereby hampering their interaction with phospholipids.

The most unexpected finding was the observation of eight NAD(H) molecules specifically bound to the dimer/dimer interface of the tetramer. Our experiments revealed the importance of NAD(H) for the stability of the tetramer and hence enzyme inhibition. Binding of NAD(H) likely involves residues from several structural elements, including the lid. Thus, NAD(H) may contribute to lid immobilization in the closed state. It is therefore hypothesized that the lid is only able to open when PlaB is present as a dimer and comes into contact with the substrate (16, 21). In other ABHs, the lid impacts substrate specificity (18), as also found here for residue S129, determining PC specificity.

NAD(H) appears to be a particularly attractive compound for controlling the activity of PlaB, since it is a central cofactor in energy homeostasis (22) expected to be confined to the intracellular milieu. Both the oxidized form NAD^+^ and the reduced form NADH inhibited PlaB. NAD^+^ is usually present in much higher amounts than NADH, and intracellular concentrations of NAD^+^ reach millimolar levels (23, 24), implying that NAD^+^ might have a greater impact on PlaB regulation.

We prefer the following model of PlaB activation and localization control: High concentrations of intracellular NAD(H) mediate tetramer formation of PlaB and thus inhibit lipolytic activity and presumably protect from bacterial lysis. Once the enzyme is exported, the concentration of NAD(H) drops significantly, leading to dissociation of the dinucleotides and consequently to separation of the tetramer into two active dimers. Dissociation of the tetramer will liberate β9/β10 from the dimer/dimer interface and thus transform the protein complex into its membrane-interacting form (Fig. 5).

**Figure 5.**
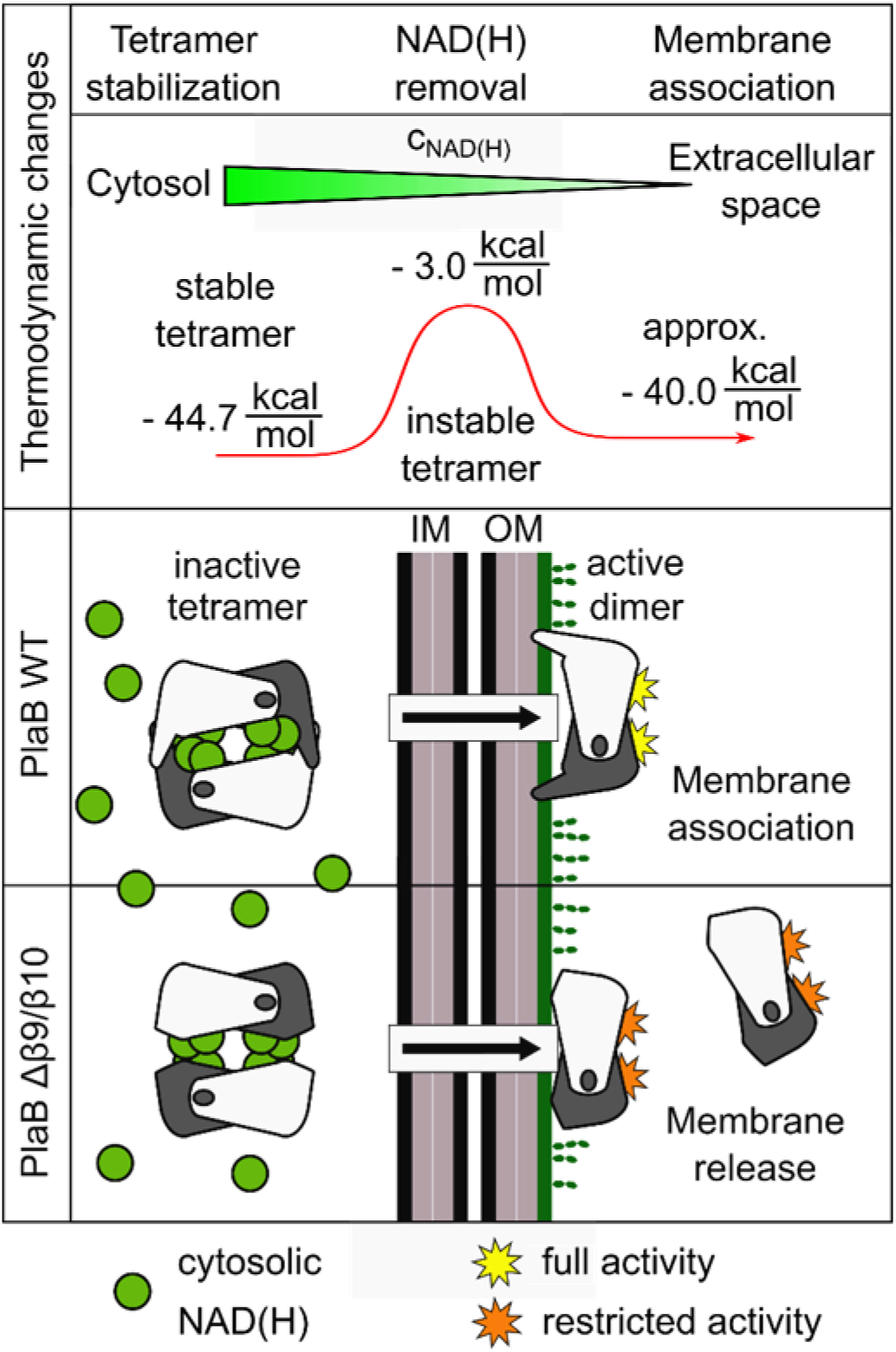
Model of NAD(H)-dependent regulation of PlaB activity and membrane association. High concentrations of intracellular NAD(H) mediate tetramer formation of PlaB and thus inhibit lipolytic activity and presumably protect from bacterial lysis. It is assumed that, upon PlaB export, NAD(H) concentrations drop and PlaB tetramers dissociate into dimers. Dimers will subsequently associate with the bacterial OM and be presented in the active form. The tetramer is stabilized by NAD(H) (ΔG = - 44.7 kcal/mol) but renders instable without the ligand (- 3.0 kcal/mol). In the dimeric form of PlaB, β9/β10 becomes accessible for membrane interaction. The expected energy gain upon membrane association of two PlaB dimers with two β9/β10 elements each can be estimated to amount to approximately - 40 kcal/mol and therefore membrane association is thermodynamically supported. Although NAD(H) still stabilizes the PlaB Δβ9/β10 tetramer, the deletion mutant is impaired in terms of OM localization and activity.

Calculations show that NAD(H) stabilizes the PlaB tetramer by - 44.7 kcal/mol whereas the nucleotide-free tetramer is unstable. Each dimer, though, is stabilized by - 29.8 kcal/mol. In the dimeric form of PlaB, β9/β10 becomes accessible for membrane interaction. The expected energy gain upon membrane association of two PlaB dimers with two β9/β10-elements each can be estimated to amount to approx. - 40 kcal/mol (25), disregarding possible further energy gain due to interactions between the CTD and the membrane. Together, this hints at a fine-tuned regulatory mechanism linking ligand-dependent oligomerization, activity and localization on the one hand for protection of *L. pneumophila* from PlaB-mediated self-lysis and on the other for presentation of the active enzyme at the host – pathogen interface.

NAD(H) acts as a cofactor of enzymes involved in numerous fundamental cellular processes (26); yet cases where NAD(H) triggers inactivity by means of tetramerization such as in PlaB have to our knowledge not yet been described. Further, several enzymes that require oligomerization to obtain full activity are known (27, 28), but the opposite, i.e. deoligomerization leading to activation, is rare (29, 30). Examples include the trimeric purine nucleoside phosphorylase (PNP) from bovine spleen, which dissociates into significantly more active monomers upon dilution (31), or the dimeric *Crotalus atrox* venom PLA_2_, which seems to split into monomers upon incubation with high concentrations of n-dodecylphosphocholine (32). However, these proteins differ in that they are not inactivated in a ligand-bound oligomeric form. Therefore, to our knowledge, PlaB is the first example of an enzyme that is inhibited by ligand-mediated oligomerization.

## Materials and Methods

### Bacterial strains and DNA techniques

Bacterial strains, plasmids, and primers are listed in Table 1. *L. pneumophila* strain Corby was used for *plaB* cloning (33). Versions of PlaB were generated by means of the QuikChange Site-Directed Mutagenesis Kit (Stratagene) (Table S4). Primers were obtained from Eurofins MWG Operon and IDT (Table S5). The bacterial strains were cultivated in lysogeny broth (LB) media containing 100 µg/mL ampicillin or 30 μg/mL chloramphenicol (for *E. coli*) or buffered yeast extract (BYE) broth when required containing 6 μg/mL chloramphenicol (for *L. pneumophila*).

### Protein expression and purification

PlaB for crystallization was produced as N-terminally Strep-tagged inactive Strep-PlaB_D203N_ as described previously (9). Briefly, protein was expressed in *E. coli* BL21 (pKK21), using 50 µM isopropyl 1-thio-D-galactopyranoside (IPTG) for induction. Seleno-*L*-methionine labeling was achieved by growing the bacteria in M9 minimal medium supplemented with 50 mg/L seleno-*L*- methionine (SeMet) and inducing with 50 µM IPTG overnight at 20 °C. Purification involved Strep-Tactin affinity and size-exclusion chromatography, followed by concentration to 10 mg/mL and storage at -80 °C until further usage. For activity and localization analyses, PlaB and variants were expressed using 0.1 mM IPTG for induction and purification by means of Strep-Tactin affinity and size-exclusion chromatography.

### Size exclusion chromatography-multiangle light scattering (SEC-MALS)

SEC-MALS experiments were performed on a chromatography system equipped with a Superose200 10 300 SEC column, a miniDAWN TREOS MALS detector and an Optilab T-rEX refractometer (Wyatt Technology Corp., USA). The column was equilibrated with 150 mM NaCl, 50 mM Tris pH 8 and 350 μl protein solution at 10 mg/mL were analyzed. Data were processed with the Astra^®^ software package (Wyatt Technology Corp., USA).

### Thermal shift assay (TSA)

TSA assays were performed by mixing 5 μL protein solution at 2 mg/mL with 5 μL SYPRO^®^Orange (1:50 diluted from 5000x stock solution, Invitrogen, USA), 5 μl of 8-fold buffer (1200 mM NaCl and 400 mM Tris pH 8), and 35 μL RUBIC Additive Screen (Molecular Dimensions, USA) in a 96-well plate. Data were collected with a C1000 Touch™ Thermal Cycler, equipped with a CFX96™ Real-Time System from 4 °C to 95 °C in 1 °C increments. TSA data was analyzed with the software BIO-RAD CFX96 Manager 3.1 (BIO-RAD, USA).

### Crystallization and data collection

Initial crystallization conditions were identified with the vapor diffusion method in sitting drops consisting of 200 nL protein solution at a concentration of 1 to 6 mg/mL mixed with the same volume of precipitant and equilibrated against 60 µL of reservoir at 20°C. Precipitants yielding crystals were optimized in grid and random screens. Microseeding experiments were performed with an OryxNano liquid dispenser (Douglas Instruments, UK), using 100 nL seed stock of native PlaB crystals broken up in 500 µL of the corresponding reservoir and 300 nl SeMet PlaB at 4 mg/mL mixed with 200 nL of precipitant solution. Crystals were further optimized by pre-incubating the protein with 1 mM thio-nicotinamidedinucleotide (SNAD). Diffraction data collection proceeded with crystals obtained with precipitants consisting of 100 mM Tris pH 8.5, 8.33 % glycerol, 10.8 % 2-propanol (native PlaB) and 77.8 mM NaSCN, 1.67 % glycerol, 5.44 % PEG400, 11.6 % 2-propanol, and 4.11 % tacsimate (SeMet PlaB). Crystals were cryoprotected in mother liquor supplemented with 10 % (v/v) (2*R*,3*R*)-(−)-2,3-butandiole before flash-cooling in liquid nitrogen. Diffraction data were collected on beamlines P11 (PETRAIII, DESY Hamburg, Germany) and PXIII (SLS, Paul Scherrer Institute, Villigen, Switzerland (34)). Indexing, integration was achieved with XDS and scaling involved AIMLESS or STARANISO (35–39), as summarized in Table S1.

### Structure determination and refinement

Initial phases were derived from single anomalous dispersion differences in diffraction data of a crystal of SeMet-labeled PlaB collected at the K-absorption edge of selenium. Heavy atom positions were identified with hkl2map (37) and forwarded to phenix.phaser of the PHENIX software suite (40). An initial model was obtained with phenix.autobuild and completed manually in Coot (41) from the CCP4 software suite (38). Further refinement involved alternating rounds of manual adjustments and optimization in phenix.refine. Model statistics are provided in Table S1. Coordinates and structure factor amplitudes have been deposited in the Protein Data Bank (PDB) (42) under accession code 6zth (seleno-*L*-methionine-containing structure with low-occupancy NAD(H)) and 6zti (unlabeled PlaB containing SNAD at full occupancy).

### Determination of lipolytic activities

Purified protein, bacterial lysates, or culture supernatants of *E. coli* BL21 expressing PlaB or its versions were incubated with dipalmitoylphosphatidylglycerol (PG), dipalmitoylphosphatidylcholine (PC), 1-monopalmitoyl-lysophosphatidylglycerol (LPG) (Avanti Polar Lipids, Inc.), and dipalmitoylphosphatidylinositol (PI) (MoBiTec). The release of fatty acids was determined using the NEFA-HR(2) assay (Wako Fujifilm) as described previously (8).

### Bacterial cell fractionation

Separation of inner and outer membranes of *E. coli* and *L. pneumophila* strains was performed with changes according to Roy & Isberg (43). For *E. coli* 100 mL of cell culture were induced at an OD_600_ = 0.8 with 0.1 mM IPTG and incubated for 3 h at 37 °C and 250 rpm. For *L. pneumophila* the induced culture was grown until late exponential phase. The culture was adjusted to OD_600_ = 1. The pellet of 100 mL was resuspended in 1 mL cold fractionation buffer (40 mM HEPES, 0.1 mM EDTA, 1x cOmplete™ Protease Inhibitor Cocktail (Roche Sigma)) and the pellet of 1 mL lysed with Bug Buster (pellet sample) (*Bug Buster® Master Mix*, Novagen® Merck). For lysis, the cells were sonified for 3 min (30% amplitude, 0.4 s pulse, Bandelin Sonoplus HD2070). The supernatant (soluble components), obtained after centrifugation at 10000 xg for 20 min at 4 °C, was ultra-centrifuged at 100;000 xg for 1 h at 4 °C. The pellet (membrane fraction) was washed and resuspended in 1 mL cold fractionation buffer and 2 % Triton X-100 and 30 mM MgCl_2_ were added (43, 44). After 5 min at room temperature, the pellets were resolved in an ultrasound bath (5 min, room temperature Bandelin Sonorex Super RK 103 H), and ultracentrifuged at 100,000 xg for 2 h at 4 °C. The supernatant (inner membrane fraction) was separated from the pellet (outer membrane), which was dissolved in 1 mL fractionation buffer using an ultrasound bath for 5 min. Fractions were analyzed by means of Western blotting detecting Strep-PlaB (StrepMAB-Classic, HRP conjugate, Iba), the outer membrane control proteins OmpA (α-OmpA was kindly provided by Prasadaro Nemani (45)) or MOMP (α-MOMP, MONOFLUO™ *Legionella pneumophila* IFA Test Kit, BioRad), the inner membrane control protein LepB (antibody kindly provided by Gunnar von Heinje (46), and the cytosolic control proteins DnaK (DnaK monoclonal antibody, Enzo Life Sciences).

### Trichloroacetic acid (TCA) protein precipitation

Detection of PlaB in culture supernatants was done using TCA precipitation and Western blotting using a Strep antibody (StrepMAB-Classic, HRP conjugate, Iba). TCA was added to a final concentration of 10 %. The sample was left for overnight incubation on ice and then centrifuged. The pellet was resuspended in ice-cold acetone three times, incubated for 30 min on ice and centrifuged. Two times more, the pellet was washed in ice-cold acetone.

## Acknowledgments

We acknowledge the staff at beamlines P11 (PETRAIII, DESY Hamburg, Germany; (47)), BL14.2 at BESSYII (Helmholtz Zentrum Berlin, Germany; (48)) and PXIII/X06DA at SLS (Paul Scherrer Institute, Villigen, Switzerland (34)) for providing us access to their facilities. We thank Dirk Heinz and Sally Pham for work in the early stages of structural analysis and Christian Galisch, Susanne Karste and Ute Strutz for excellent technical assistance. This work was supported by funds from the Deutsche Forschungsgemeinschaft awarded to AF (FL359/4-2/3) and WB (281361126/GRK2223).

