## supplemental material for "NAD(H)-mediated tetramerization controls the activity of *Legionella pneumophila* phospholipase PlaB"

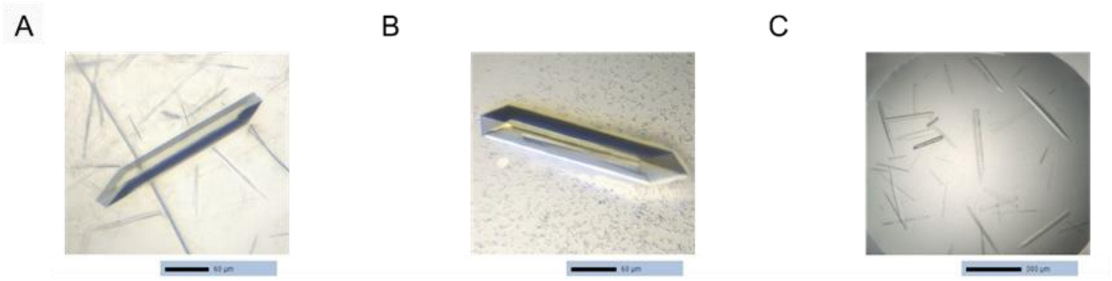


**Figure S1. PlaB crystals used in this study.**

A) PlaB crystal grown for more than 150 days. The crystal was used for the generation of a seed stock suspended in a reservoir solution.

B) Crystal grown from seleno-*L*-methionine derivatized PlaB after seeding from A). The crystal was used for SAD data collection and displayed $P2_1$ symmetry with strong tNCS.

C) Crystals grown after addition of 1 mM SNAD to the protein buffer. These crystals were used for collection of data with improved quality and displayed P1 symmetry.

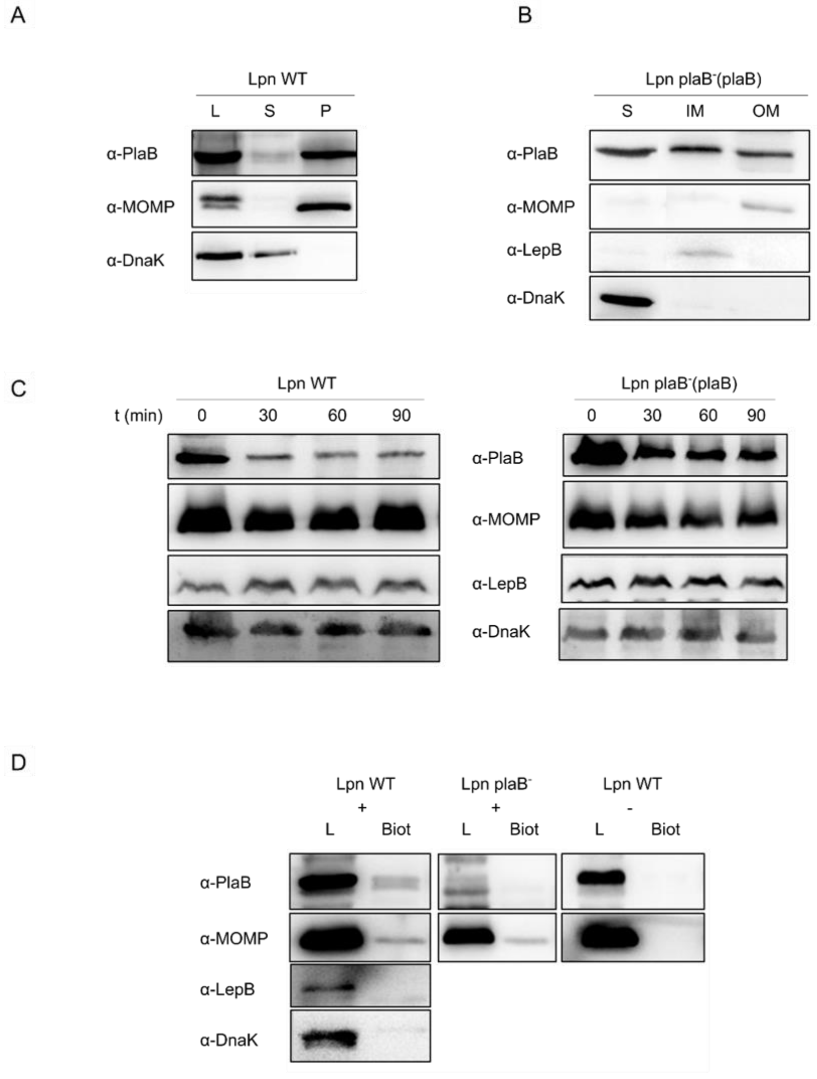


**Figure S2. PlaB is located at the outer membrane and surface of *L. pneumophila*.**

A) Cell lysate (L) of *L. pneumophila* Corby wild type (Lpn WT) was separated into cytosolic (S) and membrane-associated fractions (P) by ultracentrifugation. 20 µL of each volume-adapted fraction were analyzed by Western blotting against PlaB, MOMP, an outer membrane protein, and DnaK, a cytosolic protein.

B) The membrane-associated fraction of the complementing strain *L. pneumophila plaB*$^-$(*plaB*) expressing *plaB* [Lpn plaB$^-$ (plaB)] was further separated into a Triton X-100-soluble fraction (IM = inner membrane associated) and insoluble fraction (OM = outer membrane-associated) by a second ultracentrifugation step. 20 µL of each volume-adapted fraction were analyzed by Western blotting and α-PlaB, α-DnaK (cytosolic marker), α-LepB (inner membrane marker), and α-MOMP (outer membrane marker). Fractionation was performed according to Roy and Isberg (43).

C) 20 µL of intact *L. pneumophila* Corby wild type and complementing strain *L. pneumophila plaB*⁻(*plaB*) were analyzed by Western blotting and α-PlaB after incubation with 30µg/mL or 50 µg/mL proteinase K, respectively, at 37 °C for 0 to 90 min and inactivation with PMSF. MOMP, LepB, and DnaK served as controls. Proteinase K digests were performed according to Schunder et al. (6).

D) Intact cells of *L. pneumophila* wild type and *plaB* mutant were labeled with biotin. Biotin-labeled proteins extracted from *L. pneumophila* wild type and *plaB* mutant cell lysates (L) by using neutravidin agarose. 20 µL samples of cell lysate after biotinylation (L) and biotinylated proteins eluted from neutravidine agarose (Biot) were analyzed by Western blotting with α-PlaB, α-DnaK, α-LepB and α-DnaK. *L. pneumophila* wild type cells, which were not incubated with biotin reagent served as control. Results are representative of at least two (B) or three (A, C, D) independent experiments and are means and standard derivations from three independent experiments (C lower panel).

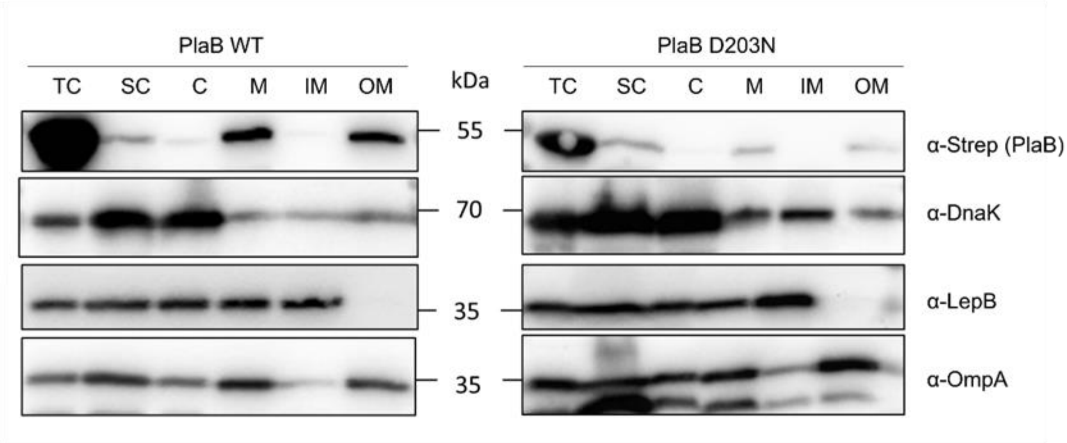


**Figure S3. Outer membrane localization of PlaB after gene expression in *E. coli.***

Western blot analysis after expression of *plaB* wild type gene or versions in *E. coli* using an anti-strep-tag antibody for PlaB detection and respective antibodies for the control proteins DnaK, LepB, and OmpA. Upper lane same experiment as shown in Fig. 2D. Abbreviations: TC – total cell lysate, SC – soluble content, C - cytosol, M – membrane, IM – inner membrane, OM – outer membrane.

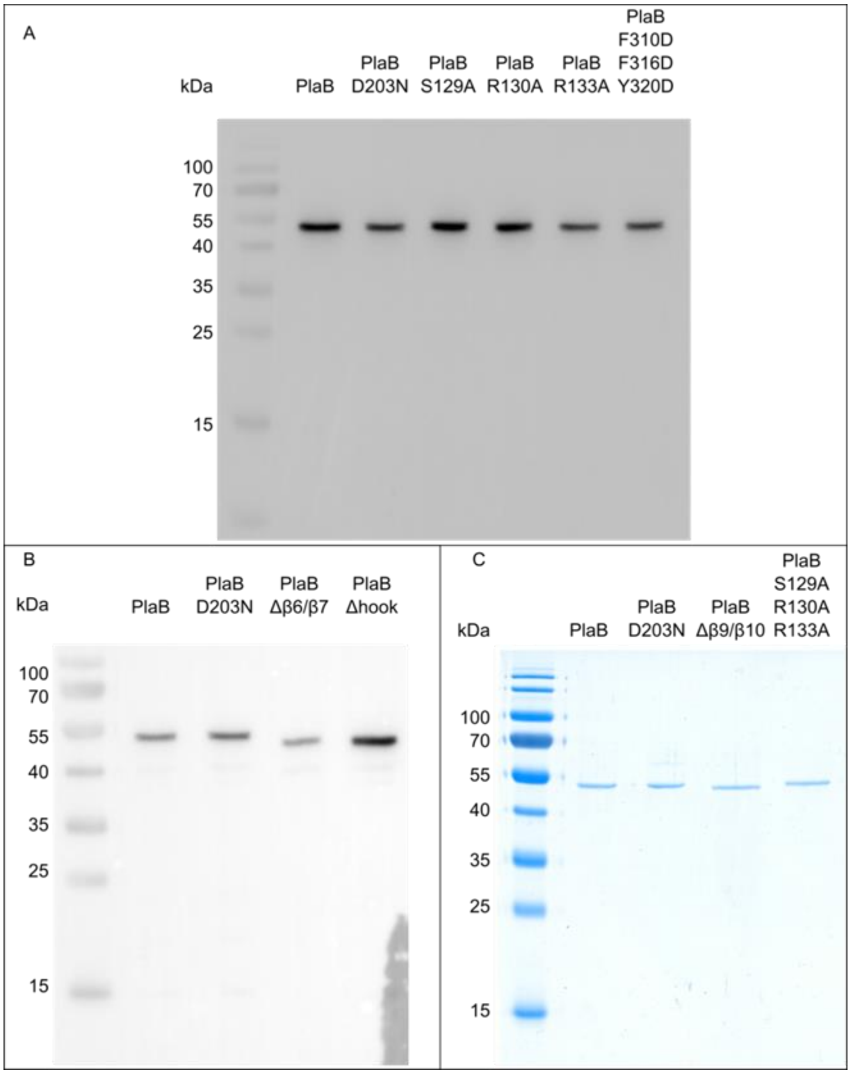


**Figure S4. Analyses of PlaB wild type and variant preparations used for enzyme activity tests.**

A and B) Western blot and SDS-PAGE of 7.5 µL cell lysate. PlaB versions are detected by using an anti-strep-tag antibody. C) SDS-PAGE of purified PlaB versions where 10 µL of a 1 µM protein solution was applied. Molecular weight markers are shown in the left lane in kDa (A-C).

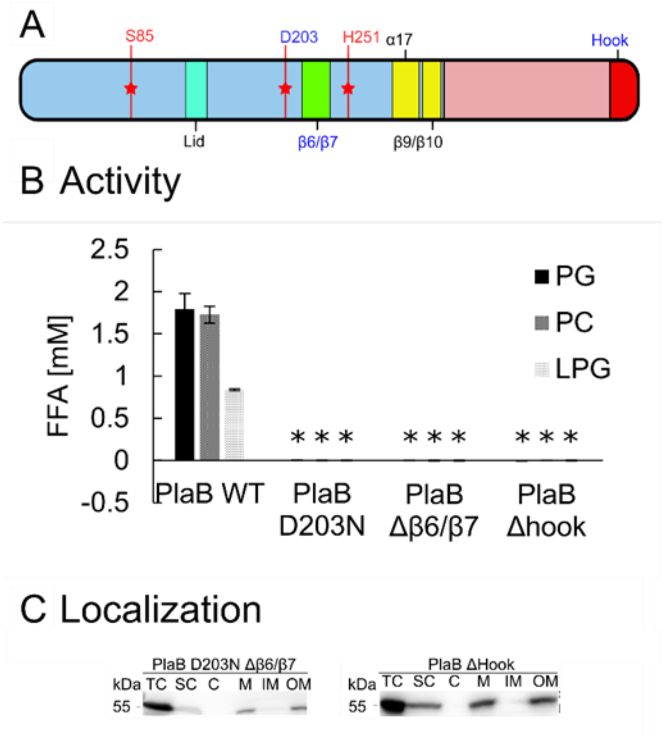


**Figure S5. PlaB β6/β7 and hook structures of PlaB are essential for activity but not localization.**

A) Overview of PlaB major sequence and structural features, such as the active site (red stars) and residues mutated in the context of this figure (blue text).

B) Enzymatic activity of PlaB WT (black) and mutant strains (D203N, Δβ6/β7, and Δhook, all no activity) towards different phospholipid substrates. 1:40 diluted cell lysates of *E. coli* expressing different PlaB versions were incubated with the different lipids for 30 min and subsequently quantities of released fatty acids (FFA) were determined. Error bars indicate SD. Statistical analysis was performed using two-tailed, unpaired Student's t-tests, relating PlaB WT to PlaB versions. * $p < 0.02$ (N=3). Western blot of cell lysates, see Fig. S4.

C) Localization of different PlaB versions. Western blot analysis after cell fractionation of *E. coli* expressing different PlaB versions using an anti-strep-tag antibody. Please refer to Fig. 2D for PlaB WT and D203N mutant analysis. Abbreviations: TC – total cell lysate, SC – soluble content, C - cytosol, M – membrane, IM – inner membrane, OM – outer membrane.

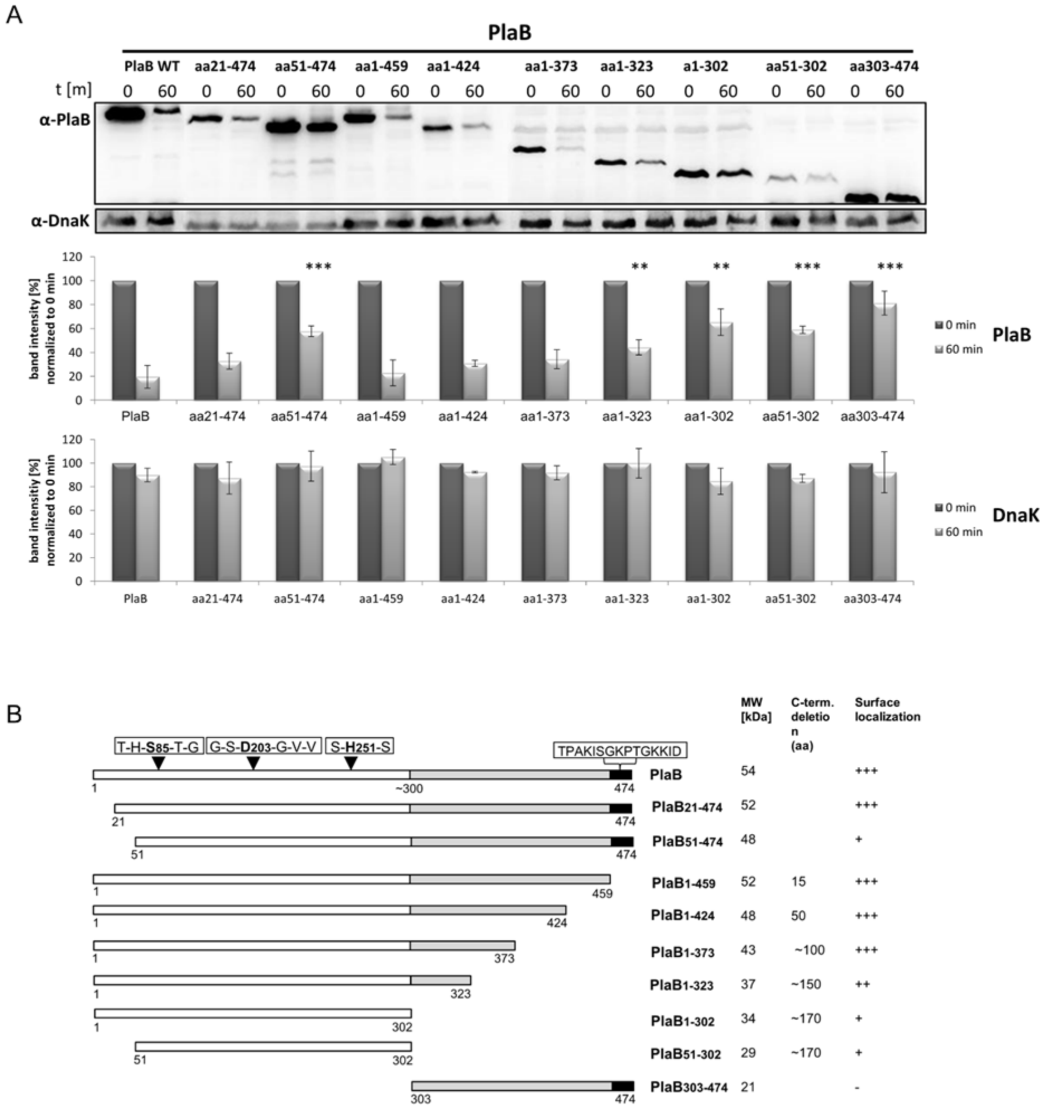


**Figure S6. Amino acids that contribute to PlaB surface presentation.**

A) Intact *L. pneumophila* Corby *plaB* mutant cells expressing C- and N-terminal truncated PlaB variants and PlaB wild type (PlaB WT) were tested for accessibility to proteinase K. 20 µL of cells were analyzed by Western blotting and α-PlaB after incubation with 50 µg/mL proteinase K at 37 °C for 60 min and inactivation with PMSF. DnaK served as control. Immunoblots of PlaB and DnaK were quantitatively analyzed by ImageJ (http://rsb.info.nih.gov/ij). Results are representative of at least three independent experiments (upper panel) and are means and standard derivations from three independent experiments (lower panel). *plaB*$^-$ strains expressing *plaB*aa51-474, *plaB*aa1-323, *plaB*aa1-302, *plaB*aa51-302 and *plaB*aa303-474 were significantly different from the wild type in all experiments, (** *P* <0.001; *** *P* <0.001, Student's *t* test, *n* 3). Proteinase K digests were performed according to Schunder et al. (6).

B) N-terminal and C-terminal truncated variants of PlaB used in the study. The catalytic triad of Ser, Asp and His (black triangles) within the N-terminal region (withe bar), the C-terminal region (grey bar), and the last 15 amino acids (black bar) are indicated. Numbers refer to amino acid positions in full-length PlaB and refer to the first and the last amino acid or the approximate start of the C-terminal region. Ability to localize at the surface is indicated.

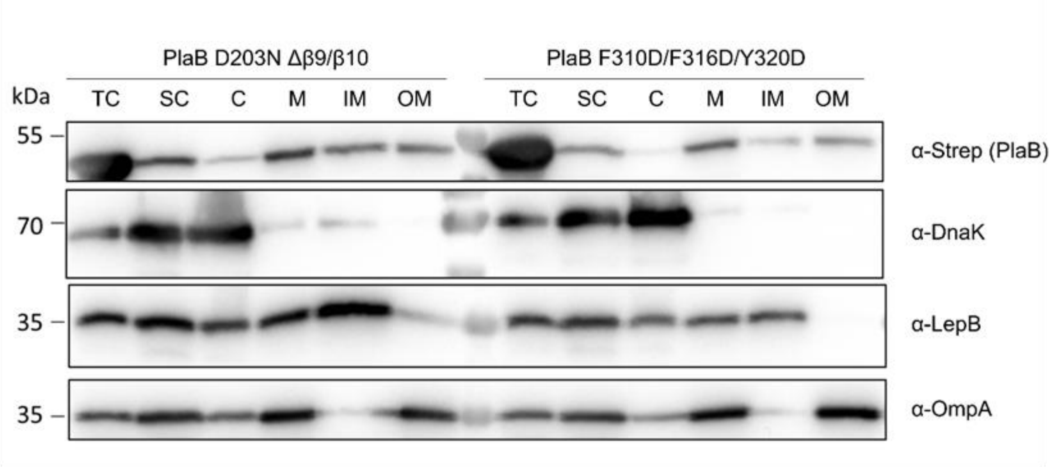


**Figure S7. β9/β10 influences localization of PlaB.**

Western blot analysis after expression of *plaB* versions in *E. coli* using an anti-strep-tag antibody for PlaB detection and respective antibodies for the control proteins DnaK, LepB, and OmpA. Please refer to Fig. 2D for PlaB WT and D203N mutant analysis. Upper lane same experiment as shown in Fig. 3D. Abbreviations: TC – total cell lysate, SC – soluble content, C - cytosol, M – membrane, IM – inner membrane, OM – outer membrane.

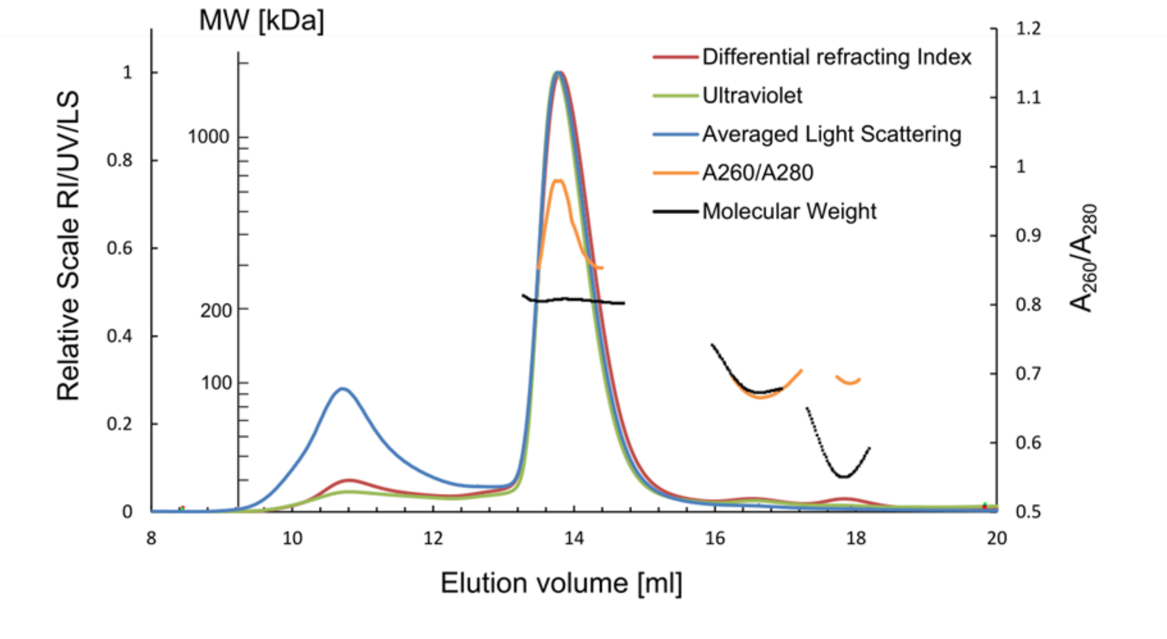


**Figure S8. Size exclusion-multiangle light scattering elution profile of PlaB**

Differential refracting index (red), ultraviolet (green) and averaged light scattering (blue) are normalized. The ratio of UV absorbance at λ = 260 nm and λ = 280 nm (yellow) and the calculated molecular weight (black) are shown for the tetramer (13.4-14.8 mL, 227.9 kDa), dimer (16.1-17.4 mL, 116.4 kDa) and monomer (17.4-18.3 mL, 60.2 kDa). At high concentrations, PlaB is present as a tetramer with a calculated mass of 228 kDa, eluting at 14 mL with an increased $A_{260}/A_{280}$ ratio, indicative of the presence of nucleotides.

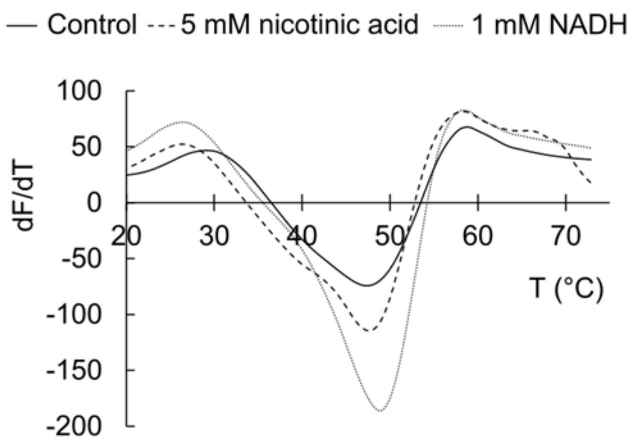


**Figure S9. Melting peaks derived from thermal shift assay on PlaB.**

PlaB without addition of any chemicals shows a flat curve with a minimum at 47°C and a shoulder at approximately 40°C. Upon the addition of nicotinic acid or NADH, the melting profile is shifted towards a more defined peak at 49°C.

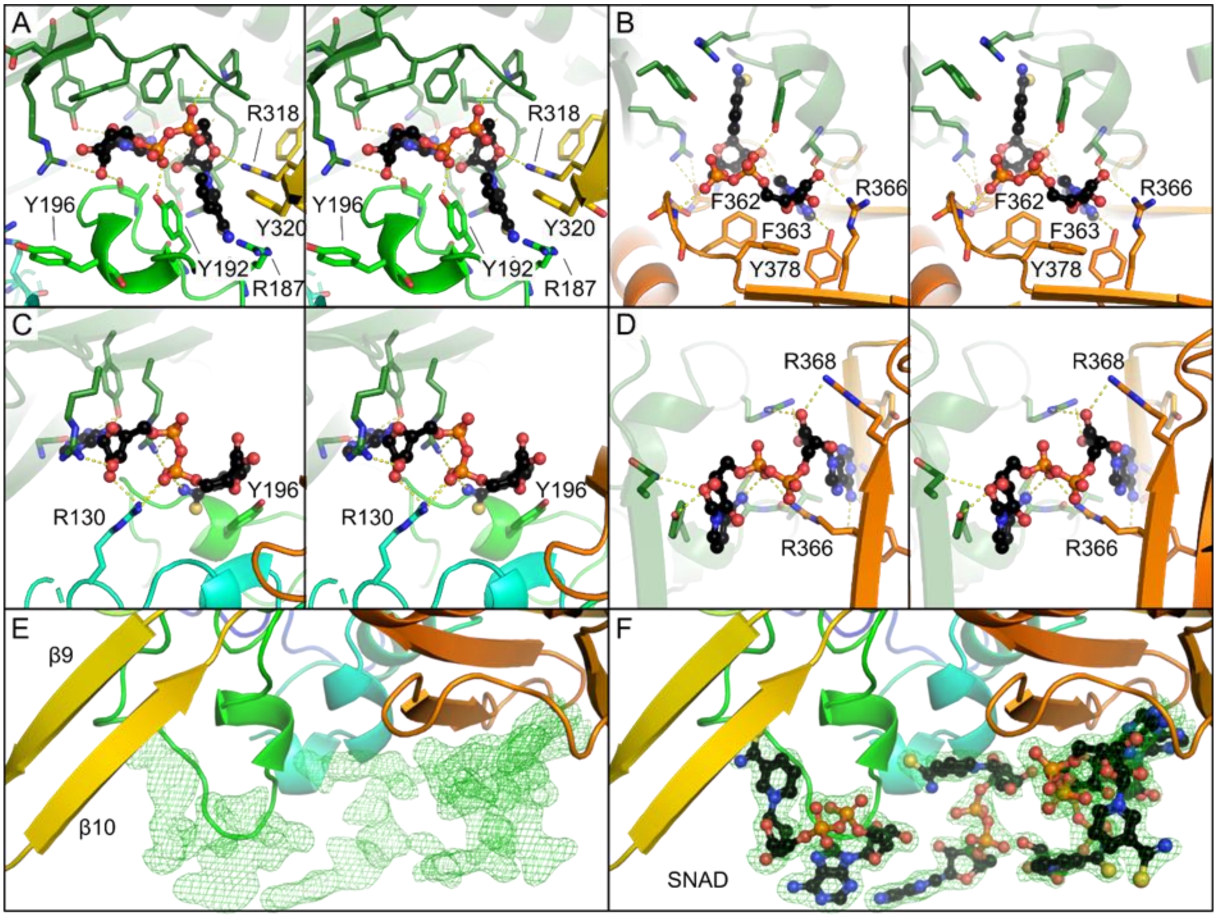


**Figure S10. PlaB binds two NAD(H) molecules per monomer.**

Panels A/B and C/D show details of the interactions within these binding sites from two different perspectives (stereo plots). Note that the identity of these ligands is SNAD, since the respective structure was obtained by cocrystallization with this ligand. Panel E and F display $|F_O - F_C|$ difference electron density of four NAD(H) binding sites at 3 σ before incorporation of SNAD into the model (green). Compare to Fig. 4A for the position of all eight NAD(H) binding sites within the PlaB tetramer.

**Table S1. Data collection and refinement statistics.**

| | **PlaB-SeMet**[a] | **PlaB/SNAD complex**[b] |
|---|---|---|
| **Data collection statistics** | | |
| Beamline | PetraIII P11 | SLS PXIII |
| Wavelength(Å) | 0.9795 | 1.21233 |
| Space group | $P2_1$ | P1 |
| Unit cell dimensions | | |
| *a*, *b*, *c* (Å) | 75.81 170.58 93.49 | 75.83 90.73 92.79 |
| α, β, γ (°) | 90.0 92.86 90.0 | 78.44 90.26 75.41 |
| Resolution range (Å) (highest shell) | 49.24 – 2.30 (2.34 - 2.30) | 46.91 – 1.81 (1.95 – 1.81) |
| Ellipsoidal resolution (Å) (direction) | - | 1.86 (0.752 **a*** - 0.142 **b*** - 0.644 **c***) |
| | - | 2.14 (0.132 **a*** + 0.971 **b*** - 0.199 **c***) |
| | - | 1.81 (0.473 **a*** + 0.360 **b*** + 0.804 **c***) |
| Total no. of reflections | 728092 (35630) | 2096239 (70139) |
| No. of unique reflections | 104870 (5140) | 161898 (8096) |
| Avg multiplicity | 6.9 (6.9) | 12.9 (8.7) |
| % Completeness (spherical) | 99.9 (99.9) | 76.0 (18.8) |
| % Completeness (ellipsoidal) | - | 91.8 (56.9) |
| $I/\sigma(I)$ | 5.9 (1.5) | 7.3 (1.6) |
| $R_{merge}$ | 0.24 (1.25) | 0.39 (2.38) |
| $R_{meas}$ | 0.26 (1.35) | 0.41 (2.60) |
| $R_{pim}$ | 0.10 (0.51) | 0.11 (0.82) |
| $CC_{½}$ | 0.99 (0.72) | 0.99 (0.53) |
| **Refinement statistics** | | |
| Resolution (Å) | 49.2 – 2.30 | 46.9 – 1.81 |
| No. of reflections used | 104627 | 161735 |
| $R_{work}$ *(%)* | 36.38 | 18.56 |
| $R_{free}$ *(%)* | 41.27 | 23.40 |
| No. of residues | | |
| protein | 1881 | 1861 |
| water | 543 | 1702 |
| ligands | 8 | 12 |
| Mean B factor ($Å^2$) | | |
| All protein residues | 38 | 21 |
| water | 26 | 30 |
| ligands | 32 | 14 |
| RMSD | | |
| Bond length (Å) | 0.006 | 0.006 |
| Bond angle (°) | 0.632 | 0.768 |
| Ramachandran plot (%) | | |
| Favored regions | 93.9 | 97.0 |
| Allowed regions | 4.9 | 3.0 |
| Outliers | 1.2 | 0 |
| **PDB code** | **6ZTH** | **6ZTI** |

[a]These data displayed strong translational non-crystallographic symmetry (tNCS; peak height relative to the origin peak: 41% at fractional coordinates 0.284, 0.500, -0.197 according to

analysis with XTRIAGE from the PHENIX software suite (40)), explaining the sub-optimal refinement statistics of the deposited structure.

[b]Statistics refer to data truncated by STARANISO to remove weak reflections affected by anisotropy (39). The program computed the indicated ellipsoid to determine anisotropic resolution limits and completeness by applying a local mean $I/\sigma(I)$ threshold of 1.5 and calculating the fraction of observed data lying inside the ellipsoid.

**Table S2. Secondary structural elements of PlaB. β-strands and α-helices are enumerated separately from element 1 to 18 and 1 to 20, respectively.** The N-terminal phospholipase domain ranges from M1 to R324 and includes most elements that form the α/β-hydrolase fold. Elements forming the α/β-hydrolase fold are marked with an *. The active site S85/D203/H251 is incorporated between corresponding secondary structure elements. The C-terminal domain ranges from Y325 to D474 and shows a bilobed β-sandwich. Sheets that form the upper lobe are marked with an $^{\dagger}$, while sheets from the lower lobe are marked with $^{\dagger\dagger}$.

| | Secondary Structure Element | Start | - | End | Assigned function | |
|---|---|---|---|---|---|---|
| N-terminal phospholipase domain (M1 – R324) | β-Strand 1* | 1 | - | 7 | | |
| | α-Helix 1* | 14 | - | 17 | | |
| | α-Helix 2* | 20 | - | 30 | | |
| | β-Strand 2* | 40 | - | 45 | Dimer interface | |
| | α-Helix 3* | 54 | - | 69 | Dimer interface | |
| | α-Helix 4* | 72 | - | 75 | | |
| | β-Strand 3* | 80 | - | 84 | | |
| | Loop β3/α5 | 85 | | | Active site S85 | |
| | α-Helix 5* | 85 | - | 98 | | |
| | α-Helix 6* | 103 | - | 105 | | |
| | β-Strand 4* | 108 | - | 114 | | |
| | α-Helix 7 | 122 | - | 125 | | |
| | α-Helix 8 | 128 | - | 131 | Tetramer interface | Lid: G127-G144 |
| | α-Helix 9 | 135 | - | 138 | | |
| | α-Helix 10 | 145 | - | 151 | | |
| | α-Helix 11* | 156 | - | 167 | | |
| | α-Helix 12* | 170 | - | 173 | | |
| | β-Strand 5* | 176 | - | 182 | | |
| | α-Helix 13 | 195 | - | 198 | Tetramer interface | |
| | Loop α13/α14 | 199 | | 207 | Active site D203 | |
| | α-Helix 14* | 208 | - | 211 | | |
| | β-Strand 6$^{\dagger}$ | 216 | - | 223 | Non-canonical β-sheet, protruding into CTD | |
| | β-Strand 7$^{\dagger}$ | 230 | - | 237 | | |
| | β-Strand 8* | 240 | - | 246 | | |
| | Loop β8/α15 | 247 | | 263 | Active site H251 | |
| | α-Helix 15 | 264 | - | 269 | | |
| | α-Helix 16* | 271 | - | 281 | | |
| | α-Helix 17 | 285 | - | 305 | Non-canonical α-helix | |
| | β-Strand 9 | 308 | - | 313 | Non-canonical β-sheet and Tetramer interface/membrane localization | |
| | β-Strand 10 | 316 | - | 321 | | |
| domain (Y325 – D474) | β-Strand 11$^{\dagger}$ | 325 | - | 334 | | |
| | β-Strand 12$^{\dagger\dagger}$ | 343 | - | 349 | Tetramer interface | |
| | β-Strand 13$^{\dagger}$ | 363 | - | 368 | Tetramer interface | |
| | β-Strand 14$^{\dagger}$ | 375 | - | 381 | | |

| | | |
|---|---|---|
| α-Helix 18 | 382 - 389 | |
| α-Helix 19 | 392 - 394 | |
| β-Strand 15†† | 398 - 404 | |
| β-Strand 16†† | 416 - 423 | |
| β-Strand 17† | 436 - 444 | |
| α-Helix 20 | 450 - 452 | Hook: R446 - D474/dimer interface |
| β-Strand 18* | 453 - 457 | complementing α/β-hydrolase fold |
| | 474 | C-terminus |

**Table S3. Size exclusion-multi angle light scattering results of PlaB WT, Δβ9/β10, and F310D/F316D/Y320D and effect of $NAD^+$ on tetramerization**. In the absence of $NAD^+$, PlaB WT concentration-dependently forms tetramers and dimers. Δβ9/β10 and F310D/F316D/Y320D were found as dimers. Upon the addition of 1 mM $NAD^+$ to the SEC buffer, exclusively tetramers were found for all proteins and concentrations.

| Sample | regular SEC buffer | | | SEC buffer + 1 mM $NAD^+$ | |
|---|---|---|---|---|---|
| | **Applied amount [µg]** | **Dimer MW (kDa)** | **Tetramer MW (kDa)** | **Applied amount [µg]** | **Tetramer MW (kDa)** |
| PlaB WT | 120 | - | 216.4 | 60 | 228.3 |
| | 60 | 117.6 | 173.3 | 30 | 225.3 |
| | 30 | 118.7 | - | 15 | 221.8 |
| | mean | - | - | mean | 225.1 ± 1.2% |
| PlaB Δβ9/β10 | 120 | 107.8 | | 60 | 217.5 |
| | 60 | 111 | | 30 | 213.6 |
| | 30 | 106.3 | | 15 | 233.7 |
| | mean | 108.4 ±1.8% | | mean | 221.6 ± 3.9% |
| PlaB F310D/ F316D/ Y320D | 120 | 110.2 | | 60 | 218.7 |
| | 60 | 113.1 | | 30 | 219.1 |
| | 30 | 112.2 | | 15 | 229.2 |
| | mean | 111.8 ± 1.1% | | mean | 222.3 ± 2.2% |

**Table S4. Strains used in the study.** Abbreviations: Ec: *E. coli*, Lpn: *L. pneumophila*, TEV: cleavage site of Tobacco Etch Virus protease, * strains shown in supporting information.

| Strain | Plasmid | Comment | Usage | References |
|---|---|---|---|---|
| Ec BL21 (pKK19) | pGP172 + Strep-*plaB* | PlaB wild type | Cloning | (9) |
| Ec BL21 (pKK21) | pGP172 + Strep-*plaB*$_{D203N}$ | PlaB catalytic mutant D203N | Crystallization | this study |
| Ec BL21 (pWM32) | pGP172 + Strep-TEV-NotI-*plaB* $_{D203N}$ | PlaB catalytic mutant D203N | Activity, localization | this study |
| Ec BL21 (pWM51) | pGP172 + Strep-TEV-NotI-*plaB* $_{D203N,\ \Delta\beta9/\beta10}$ | PlaB catalytic mutant D203N and ΔK308-I321::A = β9/β10 deletion mutant | Localization | this study |
| Ec BL21 (pWM52) | pGP172 + Strep-TEV-NotI-*plaB* | PlaB wild type | Activity, localization | this study |
| Ec BL21 (pWM56) | pGP172 + Strep-TEV-NotI-*plaB* $_{D203N,\ \Delta\beta6/\beta7}$ | PlaB catalytic mutant D203N and ΔL217-T236::A = β6/β7 deletion mutant | Localization | this study |
| Ec BL21 (pWM61) | pGP172 + Strep-TEV-NotI-*plaB*$_{\Delta\beta6/\beta7}$ | PlaB ΔL217-T236::A = β6/β7 deletion mutant | Activity | this study |
| Ec BL21 (pWM62) | pGP172 + Strep-TEV-NotI-*plaB*$_{\Delta\beta9/\beta10}$ | PlaB ΔK308-I321::A = β9/ β10 deletion mutant | Activity, localization | this study |
| Ec BL21 (pWM65) | pGP172 + Strep-TEV-NotI-*plaB*$_{F310D/F316D/Y320D}$ | PlaB β9/β10 region putative Π interaction mutant F310D/F316D/Y320D | Activity, localization | this study |
| Ec BL21 (pWM77) | pGP172 + Strep-TEV-NotI-*plaB*$_{1\text{-}445}$ | PlaB hook deletion mutant of the 28 C-terminal amino acids 446-474 | Activity, localization | this study |
| Ec BL21 (pWM80) | pGP172 + Strep-TEV-NotI- *plaB*$_{S129A}$ | PlaB lid mutant S129A | Activity | this study |
| Ec BL21 (pWM81) | pGP172 + Strep-TEV-NotI- *plaB*$_{R130A}$ | PlaB lid mutant R130A | Activity | this study |
| Ec BL21 (pWM82) | pGP172 + Strep-TEV-NotI- *plaB*$_{R133A}$ | PlaB lid mutant R133A | Activity | this study |
| Ec BL21 (pWM83) | pGP172 + Strep-TEV-NotI-*plaB*$_{S129A/R130A/R133A}$ | PlaB lid mutant S129A/R130A/R133A | Activity, localization | this study |
| Lpn Corby* | - | Wild type strain Corby | localization | (33) |
| Lpn Corby Δ*plaB* * | - | *plaB* knock out mutant | localization | (7) |
| Lpn Corby Δ*plaB* (pJB04)* | pBCKS + *plaB* | PlaB wild type | localization | (8) |
| Lpn Corby Δ*plaB* (pKK48)* | pBCKS + *plaB* $_{21\text{-}474}$ | PlaB deletion mutant of N-terminal amino acids 1-20 | localization | this study |
| Lpn Corby Δ*plaB* (pKK49)* | pBCKS + *plaB* $_{51\text{-}474}$ | PlaB deletion mutant of N-terminal amino acids 1-50 | localization | this study |
| Lpn Corby | pBCKS + *plaB* $_{1\text{-}459}$ | PlaB deletion mutant of | localization | this study |

| | | | | |
|---|---|---|---|---|
| Δ*plaB* (pKK50)* | | 15 C-terminal amino acids | | |
| Lpn Corby Δ*plaB* (pKK51)* | pBCKS + *plaB* $_{1\text{-}424}$ | PlaB deletion mutant of 50 C-terminal amino acids | localization | this study |
| Lpn Corby Δ*plaB* (pKK54)* | pBCKS + *plaB* $_{1\text{-}373}$ | PlaB deletion mutant of 101 C-terminal amino acids | localization | this study |
| Lpn Corby Δ*plaB* (pKK55)* | pBCKS + *plaB* $_{1\text{-}323}$ | PlaB deletion mutant of 151 C-terminal amino acids | localization | this study |
| Lpn Corby Δ*plaB* (pKK58)* | pBCKS + *plaB* $_{1\text{-}303}$ | PlaB deletion mutant of 171 C-terminal amino acids | localization | this study |
| Lpn Corby Δ*plaB* (pKK59)* | pBCKS + *plaB* $_{51\text{-}323}$ | PlaB deletion mutant of N-terminal amino acids 1-50 and of 151 C-terminal amino acids | localization | this study |
| Lpn Corby Δ*plaB* (pKK60)* | pBCKS + *plaB* $_{303\text{-}474}$ | PlaB C-terminal half encompassing amino acids 303-474 | localization | this study |

**Table S5. Plasmids and primers used in the study.** Abbreviations: TEV: cleavage site of Tobacco Etch Virus protease, * strains shown in supporting information.

| Plasmid | Gene | Resistance/ Template | Primer name | Sequence |
|---|---|---|---|---|
| pKK21 | pGP172 + *plaB*$_{D203N}$ | Amp/ pKK19 | strepplaBD203Nf | GAATCTGGATCCAATGGG GTGGTAC |
| | | | strepplaBD203Nr | GTACCACCCCATTGGATCC AGATTC |
| pKK48* | pBCKS + *plaB*$_{21\text{-}474}$ | Cm/pJB04 | pJB04-20Ntf | CAAGGAGCGTTATGCCTC AATGGCTTGAAAATCAG |
| | | | pJB04-20Ntr | GCCATTGAGGCATAACGC TCCTTGAATCGTTG |
| pKK49* | pBCKS + *plaB*$_{51\text{-}474}$ | Cm/pJB04 | pJB04-50Ntf | CAAGGAGCGTTATGACGG TGACGGTTGACGATATAG |
| | | | pJB04-50Ntr | CCGTCACCGTCATAACGCT CCTTGAATCGTTG |
| pKK50* | pBCKS + *plaB*$_{1\text{-}459}$ | Cm/pJB04 | pJB04-15Ctf | CAACAACCTTTGATACCTC TTTCGTTTGATGC |
| | | | pJB04-15Ctr | GAGGTATCAAAGGTTGTTG CTGATACGGAAAACTG |
| pKK51* | pBCKS + *plaB*$_{1\text{-}424}$ | Cm/pJB04 | pJB04-50Ctfbl | TGATACCTCTTTCGTTTGA TGC |
| | | | pJB04-50Ctrbl | CGATGAATGAAAATCAAGT AACC |
| pKK54* | pBCKS + *plaB*$_{1\text{-}373}$ | Cm/pJB04 | pJB04-100Ctf | GCGAAATCTCAATAATCGA TGAAAACTGACTTATTTTC |
| | | | pJB04-100Ctr | GAAAATAAGTCAGTTTTCA TCGATTATTGAGATTTCGC |
| pKK55* | pBCKS + *plaB*$_{1\text{-}323}$ | Cm/pJB04 | pJB04-150Ctf | CGAGTATATAACCAATTGA TACTCTATGATTATTTTCC |
| | | | pJB04-150Ctr | GGAAAATAATCATAGAGTA TCAATTGGTTATATACTCG |
| pKK58* | pBCKS + *plaB*$_{1\text{-}302}$ | Cm/pJB04 | pJB04-170Ctf | CCTTATGCTCATTTTTCTAA GTCTCTTTAG |
| | | | pJB04-170Ctr | CTAAAGAGACTTAGAAAAA TGAGCATAAGG |
| pKK59* | pBCKS + *plaB*$_{51\text{-}323}$ | Cm/pKK55 | pJB04-50Ntf pJB04-50Ntr/ | CAAGGAGCGTTATGACGG TGACGGTTGACGATATAG |
| | | | | CCGTCACCGTCATAACGCT CCTTGAATCGTTG |
| pKK60* | pBCKS + *plaB*$_{303\text{-}474}$ | Cm/pJB04 | Ct303474strepf | GAGCGTTATGCAGAAAAAT GAGCATAAGGAATTTG |
| | | | Ct303474strepr | CATTTTTCTGCATAACGCT CCTTGAATCGTTGTC |
| pWM32 | pGP172 + Strep-TEV-*plaB*$_{D203N}$ | Amp/ pKK21 | pKK21_TEV3_fw | TCGAAAAAGGCGCCGGTA CCGAGAACCTCTACTTCCA GGGAGGCGGCCGCATGAT TGTTA |
| | | | pKK21_TEV3_rv | TAACAATCATGCGGCCGC CTCCCTGGAAGTAGAGGT TCTCGGTACCGGCGCCTT TTTCGA |

| | | | | |
|---|---|---|---|---|
| pWM51 | pGP172 + Strep-TEV-*plaB*$_{D203N, \Delta\beta9/\beta10}$ | Amp/ pWM32 | PlaB_sheet2_fw | GCAACCAATCGTTACTCTATGATTATTTTC |
| | | | PlaB_sheet2_rv | ATGCTCATTTTTCTGAGTCTCTTTAG |
| | | | plaBsheet2_Ala_fw | AATGAGCATGCAACCAATCGTTA |
| | | | plaBsheet2_Ala_rv | TAACGATTGGTTGCATGCTCATT |
| pWM52 | pGP172 + TEV-*plaB* | Amp/ pWM32 | PlaB_revert_fw | GAATCTGGATCCGATGGGGTGGTA |
| | | | PlaB_revert_rv | TACCACCCCATCGGATCCAGATTC |
| pWM56 | pGP172 + Strep-TEV-*plaB*$_{D203N, -\Delta\beta6/\beta7}$ | Amp/ pWM32 | PlaB_sheet1_fw | GCACGAACACAGCCAATGGCATT |
| | | | PlaB_sheet1_rv | GCTATAATTCATATTTGTTGCAGCAAC |
| | | | plaBsheet1_AA216_fw | CAACAAATATGAATTATAGCGCACGAACACAGCCAATG |
| | | | plaBsheet1_AA216_rv | CATTGGCTGTGTTCGTGCGCTATAATTCATATTTGTTG |
| pWM61 | pGP172 + Strep-TEV-*plaB*$_{-\Delta\beta6/\beta7}$ | Amp/ pWM56 | PlaB_revert_fw | GAATCTGGATCCGATGGGGTGGTA |
| | | | PlaB_revert_rv | TACCACCCCATCGGATCCAGATTC |
| pWM62 | pGP172 + Strep-TEV-*plaB*$_{\Delta-\Delta\beta9/\beta10}$ | Amp/ pWM51 | PlaB_revert_fw | GAATCTGGATCCGATGGGGTGGTA |
| | | | PlaB_revert_rv | TACCACCCCATCGGATCCAGATTC |
| pWM65 | pGP172 + Strep-TEV-*plaB*$_{F310D/F316D/Y320D}$ | Amp/ pWM52 | PlaB_Y320D_fw3 | GCGAGGATATAACCAATCGTTACTCTATGATTATTTTC |
| | | | PlaB_Y320D_rv3 | GTTATATCCTCGCGCGTAAAGACAAGTGTTTTC |
| | | | PlaB_F316D_fw2 | CTTGTCGATACGCGCGAGGATATAACCAATCG |
| | | | PlaB_F316D_rv2 | GCGTATCGACAAGTGTTTTCACATCTTCCTTATG |
| | | | PlaB_F310D_fw2 | AGGAAGATGTGAAAACACTTGTCGATACGCGC |
| | | | PlaB_F310D_rv2 | TCACATCTTCCTTATGCTCATTTTTCTGAGTCTC |
| pWM77 | pGP172 + Strep-TEV-*plaB*$_{1-445}$ | Amp/ pWM52 | PlaB_Stop446_fw | GTTGAAATCATGCTTCAATGACGAGTAGATAGGAC |
| | | | PlaB_Stop446_rv | GTCCTATCTACTCGTCATTGAAGCATGATTTCAAC |
| pWM80 | pGP172 + Strep-TEV-*plaB*$_{S129A}$ | Amp/ pWM52 | PlaB_S129A_fw | CAAAGCCCGCCTAGGCCGCATAAAGAGTTTTTTTG |
| | | | PlaB_S129A_rv | CGGGCTTTGCCAAGCTGAGCAAGAGCAG |
| pWM81 | pGP172 + Strep-TEV-*plaB*$_{R130A}$ | Amp/ pWM52 | PlaB_R130A_fw | CAAATCCGCCCTAGGCCGCATAAAGAGTTTTTTTG |
| | | | PlaB_R130A_rv | CTAGGGCGGATTTGCCAAGCTGAGCAAGAG |

| | | | | |
|---|---|---|---|---|
| pWM82 | pGP172 + Strep-TEV-*plaB*$_{R133A}$ | Amp/ pWM52 | PlaB_R133A_fw | CTAGGCGCCATAAAGAGTTTTTTTGAAGGCATTG |
| | | | PlaB_R133A_rv | CTTTATGGCGCCTAGGCGGGATTTGCCAAG |
| pWM83 | pGP172 + Strep-TEV-*plaB*$_{S129A/R130A/R133A}$ | Amp/ pWM52 | PlaB_129_130_133_fw | CAAAGCCGCCCTAGGCGCCATAAAGAGTTTTTTTG |
| | | | PlaB_129_130_133_rv | CTAGGGCGGCTTTGCCAAGCTGAGCAAGAGCAG |
